# Genetic analysis of the transition from wild to domesticated cotton (*G. hirsutum* L.)

**DOI:** 10.1101/616763

**Authors:** Corrinne E. Grover, Mi-Jeong Yoo, Meng Lin, Matthew D. Murphy, David B. Harker, Robert L. Byers, Alexander E. Lipka, Guanjing Hu, Daojun Yuan, Justin L. Conover, Joshua A. Udall, Andrew H. Paterson, Michael A. Gore, Jonathan F. Wendel

**Affiliations:** Department of Ecology, Evolution and Organismal Biology, Iowa State University, Ames, IA 50011; Plant Breeding and Genetics Section, School of Integrative Plant Science, Cornell University, Ithaca, NY 14853; Department of Crop Sciences, University of Illinois, Urbana, IL 61801; Plant and Wildlife Science Department, Brigham Young University, Provo, UT 84602; Plant Genome Mapping Laboratory, University of Georgia, Athens, GA 30602

**Keywords:** QTL, domestication, *Gossypium hirsutum*, cotton

## Abstract

The evolution and domestication of cotton is of great interest from both economic and evolutionary standpoints. Although many genetic and genomic resources have been generated for cotton, the genetic underpinnings of the transition from wild to domesticated cotton remain poorly known. Here we generated an intraspecific QTL mapping population specifically targeting domesticated cotton phenotypes. We used 466 F_2_ individuals derived from an intraspecific cross between the wild *Gossypium hirsutum* var. *yucatanense* (TX2094) and the elite cultivar *G. hirsutum* cv. Acala Maxxa, in two environments, to identify 120 QTL associated with phenotypic changes under domestication. While the number of QTL recovered in each subpopulation was similar, only 22 QTL were considered coincident (i.e., shared) between the two locations, eight of which shared peak markers. Although approximately half of QTL were located in the A-subgenome, many key fiber QTL were detected in the D-subgenome, which was derived from a species with unspinnable fiber. We found that many QTL are environment-specific, with few shared between the two environments, indicating that QTL associated with *G. hirsutum* domestication are genomically clustered but environmentally labile. Possible candidate genes were recovered and are discussed in the context of the phenotype. We conclude that the evolutionary forces that shape intraspecific divergence and domestication in cotton are complex, and that phenotypic transformations likely involved multiple interacting and environmentally responsive factors.

**Summary:** An F_2_ population between wild and domesticated cotton was used to identify QTL associated with selection under domestication. Multiple traits characterizing domesticated cotton were evaluated, and candidate genes underlying QTL are described for all traits. QTL are unevenly distributed between subgenomes of the domesticated polyploid, with many fiber QTL located on the genome derived from the D parent, which does not have spinnable fiber, but a majority of QTL overall located on the A subgenome. QTL are many (120) and environmentally labile. These data, together with candidate gene analyses, suggest recruitment of many environmentally responsive factors during cotton domestication.

## Introduction

The cotton genus (*Gossypium*) represents the largest source of natural textile fiber worldwide. Although four species of cotton were independently domesticated, upland cotton (*G. hirsutum* L.) accounts for more than 90% of global cotton production. Native to the northern coast of the Yucatan peninsula in Mexico, *G. hirsutum* is now widely cultivated across the globe (Wendel and Albert 1992). Domestication of *G. hirsutum* occurred circa 5,000 years ago, producing many phenotypic changes common to plant domestication, including decreased plant stature, earlier flowering, and loss of seed dormancy. An additional primary target unique to cotton domestication was the single-celled epidermal trichomes (i.e., fibers) that cover the cotton seed. Cotton fiber morphology varies greatly in length, color, strength, and density among the myriad accessions that span the wild-to-domesticate continuum. As a species, *G. hirsutum* is highly diverse, both morphologically and ecologically, and has a correspondingly long and complex taxonomic history (Fryxell 1968, 1976, 1979, 1992) that includes the modern, cryptic inclusion of at least two distinct species (Wendel and Grover 2015; Gallagher *et al.* 2017). Truly wild forms of *G. hirsutum* (race *yucatanense*) occur as scattered populations in coastal regions of the semiarid tropical and subtropical zones of the Caribbean, northern South America, and Mesoamerica (Coppens d’Eeckenbrugge and Lacape 2014). These are distinguished from domesticated and feral forms by their short, coarse, brown fibers, as well as their sprawling growth habit, photoperiod sensitivity, and seed dormancy requirements, among others (Figure 1). Results from molecular marker analyses, including allozymes (Wendel and Albert 1992), restriction fragment length polymorphisms (RFLPs) (Brubaker and Wendel 1994), simple sequence repeats (SSRs) (Liu and Wendel 2002; Zhang *et al.* 2011; Tyagi *et al.* 2014; Zhao *et al.* 2015; Kaur *et al.* 2017; McCarty *et al.* 2018), SNP arrays (Hinze *et al.* 2017; Cai *et al.* 2017; Ai *et al.* 2017), and next-generation sequencing (Reddy *et al.* 2017; Fang *et al.* 2017c; Ma *et al.* 2018) have quantified genetic diversity and aspects of population structure among wild, feral, and domesticated stocks of the species, as well as the allopolyploid origin of the species. Notably, the allopolyploid origin of *G. hirsutum* includes a diploid species with no spinnable fiber, i.e., the paternal parent derived from the fiberless Mesoamerican “D-genome” clade. The maternal progenitor of the allopolyploid lineage is derived from the African “A-genome” whose two extant species have been independently domesticated for fiber production.

**Figure 1.**
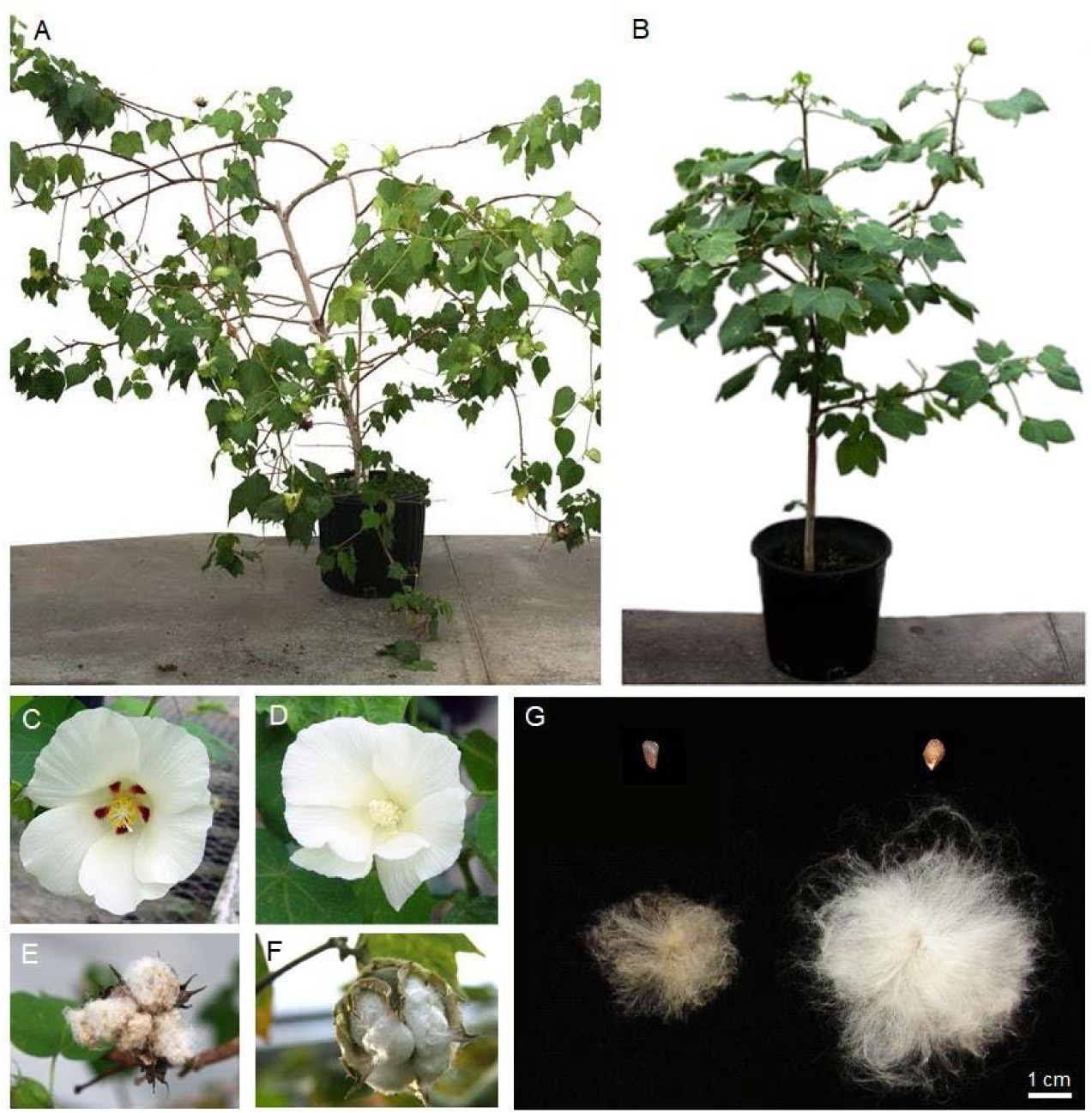
Morphological differentiation between *G. hirsutum* var. *yucatanense* TX2094 and *G. hirsutum* cv. Acala Maxxa. (A) Adult plant of TX2094, wild; (B) Adult plant of Acala Maxxa, domesticated; (C) TX2094 flower; (D) Acala Maxxa flower; (E) Open boll of TX2094; (F) Open boll of Acala Maxxa; (G) Ginned seed of TX2094 (top left) and Acala Maxxa (top right), and fiber of TX2094 (bottom left) and Acala Maxxa (bottom right). Photo credit: Kara Grupp & Mi-Jeong Yoo

Recent advances have improved our understanding of the genetic changes targeted by humans during the several millennia of cotton domestication and improvement by evaluating gene expression differences that distinguish wild and domesticated cotton fiber, either globally or for a few key genes among accessions (Haigler *et al.* 2009; Bao *et al.* 2011; Kim *et al.* 2012; Argiriou *et al.* 2012; Tuttle *et al.* 2015). Genome-scale surveys have elucidated many of the genes that are differentially expressed between wild and domesticated cotton (Hovav *et al.* 2008b; Chaudhary *et al.* 2009; Rapp *et al.* 2010; Yoo and Wendel 2014; Nigam *et al.* 2014), or among developmental stages of fiber development (Shi *et al.* 2006; Gou *et al.* 2007; Taliercio and Boykin 2007; Hovav *et al.* 2008c, 2008b; Al-Ghazi *et al.* 2009; Rapp *et al.* 2010; Wang *et al.* 2010; Yoo and Wendel 2014; Nigam *et al.* 2014; Tuttle *et al.* 2015). These many studies indicate that domestication has dramatically altered the transcriptome of cotton fiber development, but to date the specific upstream variants and interacting partners responsible for these downstream developmental differences remain to be discovered.

From a genetic perspective, multiple independent quantitative trait loci (QTL) analyses have been performed to identify chromosomal regions contributing to phenotypic variation among various cotton genotypes. Most QTL analyses to date have focused either on crosses between modern cultivars of *G. hirsutum* or on crosses between cultivated forms of *G. hirsutum* with *G. barbadense*, another cultivated species which possesses superior fiber quality but with the limitations of lower yield and a narrower range of adaptation (Fang *et al.* 2017c; Chandnani *et al.* 2017; Hu *et al.* 2019). Interspecific cotton crosses often generate negative genetic correlations between fiber quality and lint yield, and these frequently suffer from F_2_ breakdown (reviewed in (Zhang *et al.* 2014)). Taken together, these numerous studies have reported more than 2,274 QTL (Said *et al.* 2015a) pertaining to agronomically and economically important traits (e.g., plant architecture; biotic and abiotic stress resistance; fiber, boll, and seed quality and productivity). Several meta-analyses have attempted to identify possible QTL clusters and hotspots by uniting these QTL studies through a consensus map (Rong *et al.* 2007; Lacape *et al.* 2010; Said *et al.* 2015b, 2015a); QTL clusters denote genomic regions containing myriad QTL, whereas QTL hotspots are clusters of QTL for a single trait (Said *et al.* 2015b). These meta-analyses compiled QTL studies of both intraspecific *G. hirsutum* populations and interspecific *G. hirsutum* × *G. barbadense* populations, ultimately creating a QTL database from intraspecific and interspecific populations (Said *et al.* 2015a). To date, QTL analyses have yielded multiple, sometimes conflicting, insights that are accession- or environment-dependent. Some aspects of fiber development, for example, are associated with QTL enrichment in the D-subgenome of polyploid cotton (Jiang *et al.* 1998; Lacape *et al.* 2005; Han *et al.* 2006; Rong *et al.* 2007; Qin *et al.* 2008; Said *et al.* 2015b), which derives from a short fibered ancestor, but not all mapping populations reflect this bias (Ulloa *et al.* 2005; Lacape *et al.* 2010; Li *et al.* 2013). Likewise, QTL found in some environments and/or populations are not significant in similar, but non-identical, environments or in other mapping populations (Lacape *et al.* 2010; Said *et al.* 2015b, 2015a). Some data suggests that cotton fiber QTL are genomically clustered, yet with heterogeneous phenotypic effects (Rong *et al.* 2007; Qin *et al.* 2008; Lacape *et al.* 2010). Said et al. (Said *et al.* 2013, 2015b) showed that just as QTL clusters and hotspots exist for fiber quality, they also exist for other traits (e.g., yield, seed quality, leaf morphology, disease resistance), and these hotspots, while found on every chromosome, tend to concentrate in specific regions of the genome. In particular, comparisons between intraspecific and interspecific populations reveal common QTL clusters and hotspots, possibly indicative of shared genetic architecture among cultivars and between species (Said *et al.* 2015b). While these QTL analyses have increased our understanding of the number and location of chromosomal regions that contribute to differences between cultivars and species, there remains a significant gap in our understanding of genes targeted during the initial domestication of cotton and their effects, which ultimately led to the development of modern cultivars.

Here we provide an evolutionary quantitative genetics perspective on the domestication of the dominant cultivated cotton species, *G. hirsutum*, through identification and characterization of QTL for traits that have played important roles during domestication. In contrast to previous studies, we utilize an *intraspecific* cross between a truly wild form of *G. hirsutum* (var. *yucatanense*, accession TX2094) and an elite cultivar (*G. hirsutum* cv. Acala Maxxa), to bracket the “before” and “after” phenotypic characteristics of the domestication process that played out over the last 5,000 years or so. Numerous domestication-related traits were characterized in both the parents and their segregating progeny in two environments, representing characters from several broader phenotypic categories: (1) plant architecture, (2) fruiting habit, (3) phenology, (4) flower, (5) seed, (6) fiber-length, (7) fiber quality, and (8) fiber color. We generated a SNP-based genetic linkage map to anchor each QTL to the *G. hirsutum* cotton reference genome (elite accession TM1; (CottonGen; Saski *et al.* 2017)) and identify plausible candidate genes for each trait. We show that the QTL associated with *G. hirsutum* domestication are both clustered and environmentally labile. Possible candidate genes were recovered and discussed for each trait. This study provides valuable insights into the genetic basis of cotton domestication and provides information that will assist in identifying cotton domestication genes and their functional effects on cotton biology.

## Materials and Methods

### Plant Materials and Phenotyping

A total of 466 F_2_ individuals were derived from a cross between *Gossypium hirsutum* var. *yucatanense* accession TX2094 as the maternal parent (USDA GRIN accession PI 501501, collected by J. McD. Stewart) and the modern elite cultivar *G. hirsutum* cv. Acala Maxxa as the paternal parent. The *G. hirsutum* var. *yucatanense* accession was previously identified as being truly wild using both allozyme (Wendel and Albert 1992) and RFLP analysis (Brubaker and Wendel 1994), as well as by morphological evidence. To allow for the replication of alleles over time and space, these individuals were grown as two subpopulations (October 2009 to July 2010), with 232 plants located in a greenhouse at Iowa State University (Ames, Iowa), and the remaining 234 in a greenhouse at the U. S. Arid-Land Agricultural Research Center (Maricopa, Arizona); nine representatives of each parental accession were also grown in each greenhouse. At Iowa State, individual seeds were separately planted in 7.6 L (two gallon) containers containing 15:7:3:3 soil:sand:peat:perlite. Plants were grown under natural sunlight (10-11 hours of daylight) with daytime and nighttime temperatures of 25±2 and 20±2°C, respectively. Plants were fertilized twice a week with 125 ppm N. In Arizona, individual seeds were separately planted into 18.9 L (five gallon) pots containing moistened Sunshine Mix #1 (Sun Gro Horticulture Inc., Bellevue, WA) and perlite (4:1 ratio). Plants were grown under natural sunlight in a greenhouse with daytime and nighttime temperatures at 30±2 and 22±2°C, respectively. All Arizona, plants were fertilized every two-weeks with 20–20–20 (200 ppm N) Peters Professional plant nutrient solution. These two populations were subsequently evaluated for multiple traits in each of the following eight categories: (1) plant architecture, (2) fruiting habit, (3) phenology, (4) flower, (5) seed, (6) fiber length, (7) fiber quality, and (8) fiber color (Table 1). Traits were selected to cover the range of possible domestication phenotypes.

**Table 1.**
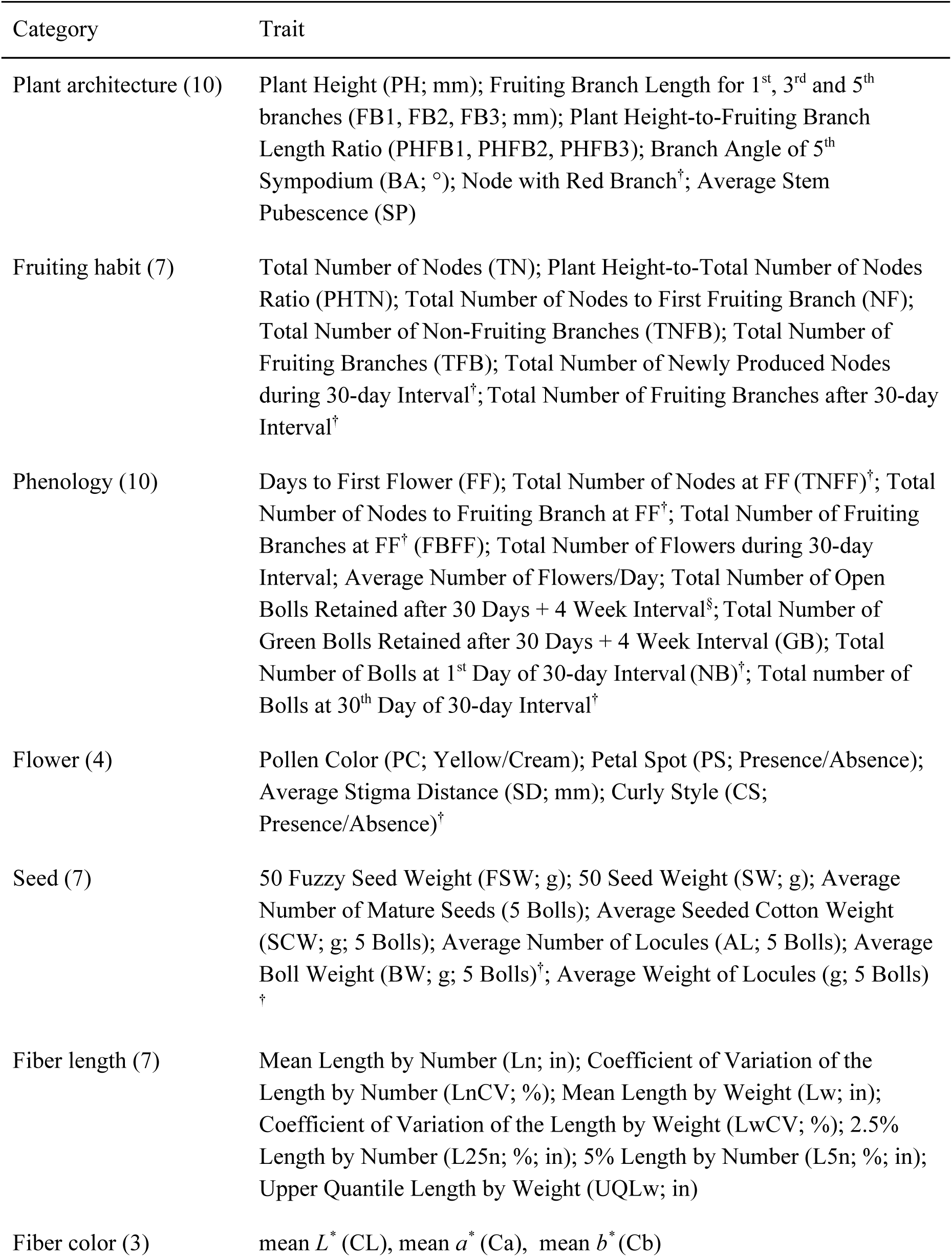
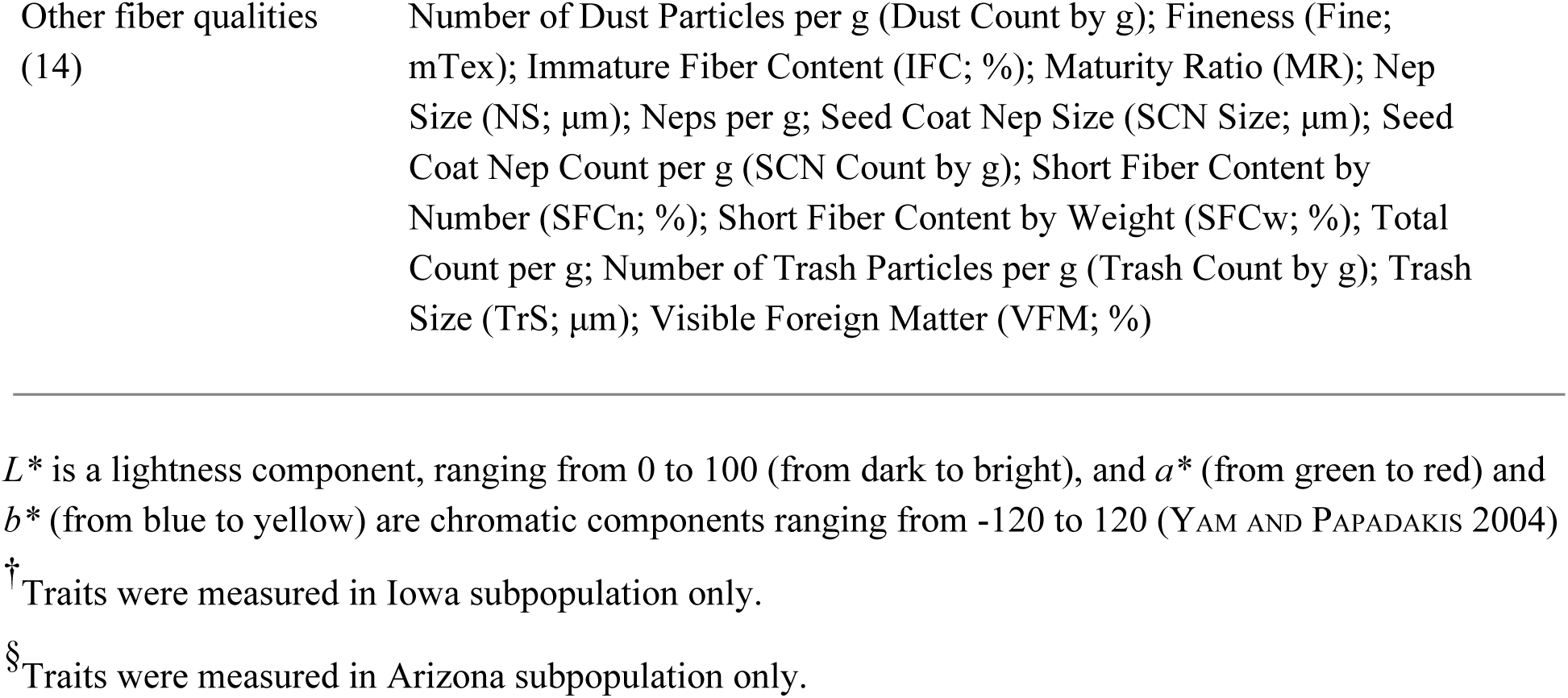
List of domestication-related traits measured in this study. For detailed information on identified QTL, refer to Table 2.

| Category | Trait |
| --- | --- |
| Plant architecture (10) | Plant Height (PH; mm); Fruiting Branch Length for 1<sup>st</sup>, 3<sup>rd</sup> and 5<sup>th</sup> branches (FB1, FB2, FB3; mm); Plant Height-to-Fruiting Branch Length Ratio (PHFB1, PHFB2, PHFB3); Branch Angle of 5<sup>th</sup> Sympodium (BA; °); Node with Red Branch<sup>†</sup>; Average Stem Pubescence (SP) |
| Fruiting habit (7) | Total Number of Nodes (TN); Plant Height-to-Total Number of Nodes Ratio (PHTN); Total Number of Nodes to First Fruiting Branch (NF); Total Number of Non-Fruiting Branches (TNFB); Total Number of Fruiting Branches (TFB); Total Number of Newly Produced Nodes during 30-day Interval<sup>†</sup>; Total Number of Fruiting Branches after 30-day Interval<sup>†</sup> |
| Phenology (10) | Days to First Flower (FF); Total Number of Nodes at FF (TNFF)<sup>†</sup>; Total Number of Nodes to Fruiting Branch at FF<sup>†</sup>; Total Number of Fruiting Branches at FF<sup>†</sup> (FBFF); Total Number of Flowers during 30-day Interval; Average Number of Flowers/Day; Total Number of Open Bolls Retained after 30 Days + 4 Week Interval<sup>§</sup>; Total Number of Green Bolls Retained after 30 Days + 4 Week Interval (GB); Total Number of Bolls at 1<sup>st</sup> Day of 30-day Interval (NB)<sup>†</sup>; Total number of Bolls at 30<sup>th</sup> Day of 30-day Interval<sup>†</sup> |
| Flower (4) | Pollen Color (PC; Yellow/Cream); Petal Spot (PS; Presence/Absence); Average Stigma Distance (SD; mm); Curly Style (CS; Presence/Absence)<sup>†</sup> |
| Seed (7) | 50 Fuzzy Seed Weight (FSW; g); 50 Seed Weight (SW; g); Average Number of Mature Seeds (5 Bolls); Average Seeded Cotton Weight (SCW; g; 5 Bolls); Average Number of Locules (AL; 5 Bolls); Average Boll Weight (BW; g; 5 Bolls)<sup>†</sup>; Average Weight of Locules (g; 5 Bolls)<sup>†</sup> |
| Fiber length (7) | Mean Length by Number (Ln; in); Coefficient of Variation of the Length by Number (LnCV; %); Mean Length by Weight (Lw; in); Coefficient of Variation of the Length by Weight (LwCV; %); 2.5% Length by Number (L25n; %, in); 5% Length by Number (L5n; %, in); Upper Quantile Length by Weight (UQLw; in) |
| Fiber color (3) | mean $L^*$ (CL), mean $a^*$ (Ca), mean $b^*$ (Cb) |
| Other fiber qualities<br>(14) | Number of Dust Particles per g (Dust Count by g); Fineness (Fine; mTex); Immature Fiber Content (IFC; %); Maturity Ratio (MR); Nep Size (NS; $\mu\text{m}$); Neps per g; Seed Coat Nep Size (SCN Size; $\mu\text{m}$); Seed Coat Nep Count per g (SCN Count by g); Short Fiber Content by Number (SFCn; %); Short Fiber Content by Weight (SFCw; %); Total Count per g; Number of Trash Particles per g (Trash Count by g); Trash Size (TrS; $\mu\text{m}$); Visible Foreign Matter (VFM; %) |

$L^*$ is a lightness component, ranging from 0 to 100 (from dark to bright), and $a^*$ (from green to red) and $b^*$ (from blue to yellow) are chromatic components ranging from -120 to 120 (YAM AND PAPADAKIS 2004)
<sup>†</sup> Traits were measured in Iowa subpopulation only.
<sup>§</sup> Traits were measured in Arizona subpopulation only.

At 150 (±7) days after planting, 10 plant architecture traits were evaluated, which include plant height, fruiting branch length, branch angle, and stem pubescence (Table 1). Data were collected for branch angles at the intersection of 1^st^, 3^rd^ and 5^th^ sympodia (secondary axes) with the main stem; however, due to high variation in the data observed from the 1^st^ and 3^rd^ sympodia, only data from the 5^th^ sympodium was considered further. In addition, the first node having a branch with red coloring was recorded in the Iowa population only (Table 1). Stem pubescence was scored independently by two people using the five-grade (1–5) ordinal scale developed by Lee (1968) (Lee 1968), where 1 is fully pubescent; the average of the two scores was recorded.

Traits relating to phenology, flowering, and fruiting were also examined. Eleven phenological traits (Table 1) were recorded, and, for consistency between the two greenhouse subpopulations, we hand-pollinated flowers for 30 days following the emergence of the first flower. Four floral traits were examined, including pollen color, the presence or absence of petal spot, average stigma distance (mm), and the presence or absence of curly styles. For pollen color, there exists a gradient of color from cream to yellow; however, we restricted our classifications to the parental color codes, i.e., “cream” vs. “yellow” observed in Acala Maxxa and TX2094, respectively. Upon maturation, seven traits related to boll/seed development were also measured on harvested bolls, such as number of mature seeds, fuzzy seed weight, and average seeded cotton weight (Table 1).

Finally, 358 fiber samples harvested from the 466 F_2_ plants were collected and sent to the Cotton Incorporated Textile Services Laboratory (Cotton Incorporated, Cary, NC) for analysis by the AFIS Pro system (Uster Technologies, Charlotte, NC), an industry standard for evaluating fiber length and other quality traits (Table 1). Fiber color was determined by a MiniScan XE Plus colorimeter (ver. 6.4, Hunter Associates Laboratory, Inc., Reston, VA), which measures color properties of *L\**, *a\**, and *b\**. *L\** is a lightness component, ranging from 0 to 100 (from dark to bright), while *a\** (from green to red) and *b\** (from blue to yellow) are chromatic components ranging from −120 to 120 (Yam and Papadakis 2004). Values were measured three times on the same fiber sample and averaged for each trait (i.e., mean *L\**, mean *a\**, and mean *b\**).

### Genotyping and Genetic Map Construction

A total of 384 KASPar-based SNP assays (277 co-dominant) were used to genotype the 466 F_2_ plants with phenotypic data (KBioscience Ltd., Hoddesdon, UK). SNP assays were designed as previously reported for *G. hirsutum* (Byers *et al.* 2012). Genomic DNA was extracted from leaf tissue using the Qiagen DNeasy Plant Mini Kit (Qiagen, Stanford, CA, USA) and normalized to an approximate concentration of 60 ng/µL.

Specific target amplification (STA) PCR was used to pre-amplify the target region of genomic DNA containing the SNPs of interest, but without the discriminating SNP base in the primer sequence. The PCR conditions for this protocol included a 15-min denaturing period at 95°C followed by 14 two-step cycles: 15 s at 95°C followed by 4 min at 60°C. This effectively increased the concentration of the target DNA relative to the remaining DNA. The sample amplicons produced by the STA protocol were then genotyped using the Fluidigm 96.96 Dynamic Arrays genotyping EP1 System (San Francisco, CA). Each Fluidigm plate run included eight control samples: two Acala Maxxa, two TX2094, two pooled parental DNA (synthetic heterozygotes), and two no-template controls (NTC). These controls served as guideposts during the genotyping process. The STA amplicons and the SNP assays were loaded onto a Fluidigm 96.96 chip, where a touchdown PCR protocol on the Fluidigm FC1 thermal cycler (San Francisco, CA, USA) was used to allow the competing KASPar primers to amplify the appropriate SNP allele in each sample.

Fluorescence intensity for each sample was measured with the EP1 reader (Fluidigm Corp, San Francisco, CA) and plotted on two axes. Some assays required more amplification in order to produce distinct clusters. For those that did not form distinct clusters during the initial analysis, an additional five cycles of PCR were performed on the plate and fluorescence intensity measured again until all assays produced sufficient resolution for cluster calling. Genotypic calls based on EP1 measurements were made using the Fluidigm SNP Genotyping Analysis program (Fluidigm 2011). All genotype calls were manually checked for accuracy and ambiguous data points that either failed to amplify and/or cluster near parental controls were scored as missing data. The final raw output for an individual chip included data from each of the multiple scans performed to ensure that the optimal amplification conditions for each assay was represented. The text output from genotyping was arranged to a compatible format for genetic mapping using Excel. Files are available at https://github.com/Wendellab/QTL_TxMx.

A genetic linkage map based on the KASPar genotyping data was constructed separately for each subpopulation using regression mapping as implemented in JoinMap4 (VAN Ooijen 2011). A LOD threshold of 5.0 was used and linkage distances were corrected with the Kosambi mapping function. Loci were excluded from the map if they failed to meet a Chi-Square test (*α* = 0.05) for expected Mendelian ratios. Separate linkage maps (i.e., not a single composite linkage map) were used for QTL analysis in each subpopulation to maximize independence when comparing results between Iowa and Arizona.

### QTL analysis

For each location, the raw phenotypic values of each trait were evaluated for statistical outliers in SAS version 9.3 (SAS Institute 2012) by examination of Studentized deleted residuals (Kutner *et al.* 2004), which were obtained from a simple linear model fitted with fixed effects for the grand mean and a single randomly sampled, representative SNP marker. QTL were detected within each greenhouse environment (Ames, IA and Maricopa, AZ) with Windows QTL Cartographer V2.5 (Wang *et al.* 2012) using the composite interval mapping (CIM) method (Zeng 1993, 1994) with a window size of 10 cM and a 1 cM walk speed. The LOD thresholds used to identify QTL were determined using a permutation test (1000 repetitions, *α* = 0.05) (Churchill and Doerge 1994), and the confidence intervals were set as the map interval corresponding to one-LOD interval on either side of the LOD peak (Mangin *et al.* 1994). If the QTL were separated by a minimum distance of 20 cM, they were considered two different QTL (Ungerer *et al.* 2002). To identify coincident QTL between subpopulations for each trait, we determined whether SNP markers were shared between QTL intervals. If at least one marker was shared between QTL marker intervals, then we concluded that the same QTL (i.e., coincident QTL) was identified in both subpopulations. A QTL cluster was declared where three or more QTL of different trait categories occurred within a 20 cM region, and a QTL hotspot was declared where three or more QTL of the same trait category occurred within a 20 cM region following (Said et al. 2015) with modification for a single genetic cross. Both QTL clusters and QTL hotspots were declared within each subpopulation, but coincident QTL clusters and QTL hotspots between subpopulations were only counted once with respect to the total of each QTL class. The linkage map showing the location of QTL (Figure 2) was generated by MapChart 2.2 (Voorrips 2002) and colorized in Adobe Photoshop Creative Suite 5 (Adobe). QTL nomenclature follows a method used in rice (McCouch *et al.* 1997), which starts with “q”, followed by an abbreviation of the trait name. The population from which the QTL derived is abbreviated at the end as “AZ” and “IA”, for Arizona and Iowa, respectively.

**Figure 2.**
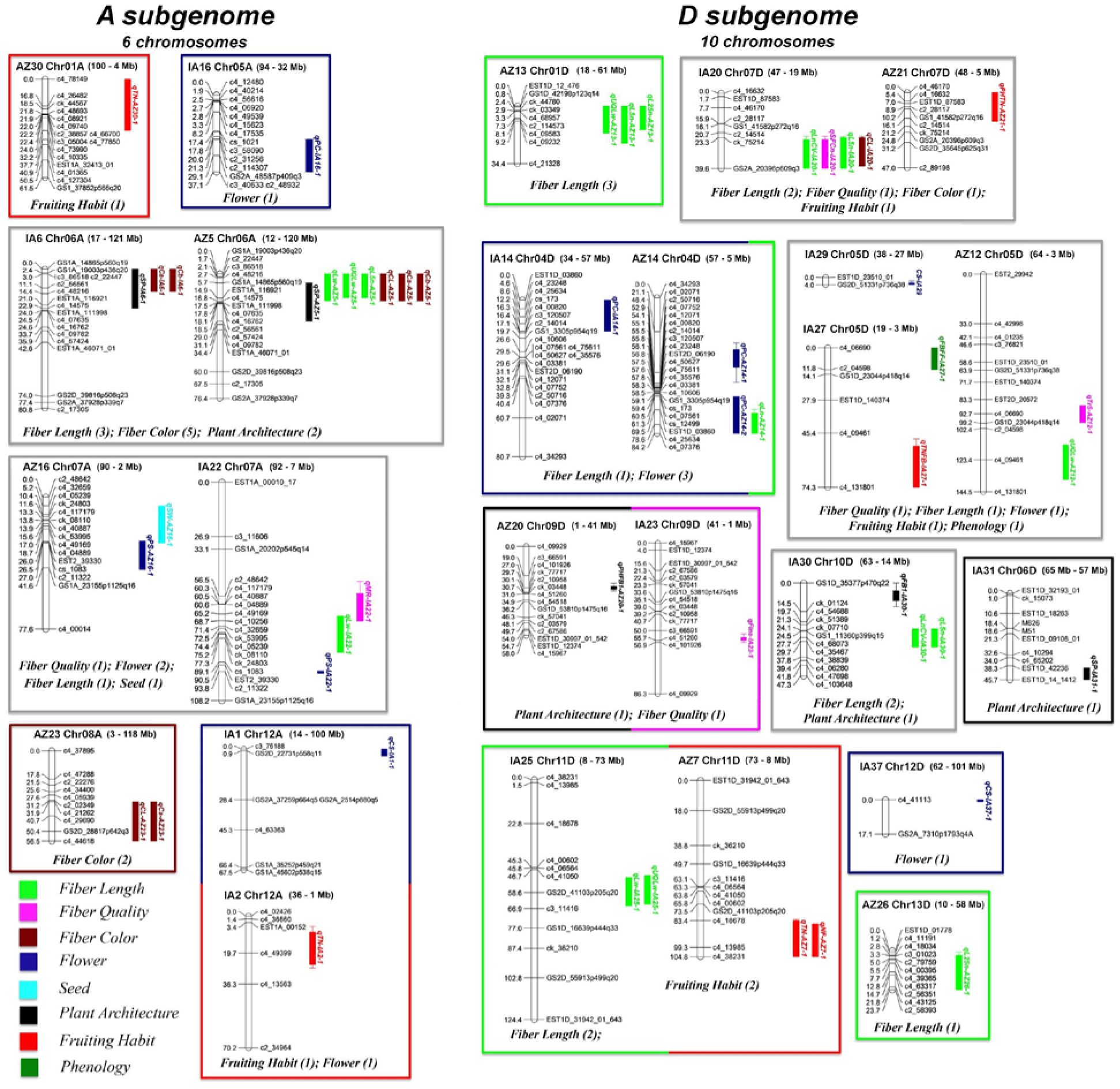
Genetic linkage map that includes the top 50 QTL associated with cotton domestication traits evaluated here, as generated by MapChart 2.2 (Voorrips 2002). While all chromosomes were recovered for the linkage map, only those linkage groups/chromosomes containing QTL are depicted here. QTL nomenclature follows that first used in rice (McCouch *et al.* 1997), which starts with “q”, followed by an abbreviation of the trait name. Environments are designated at the end of the QTL name with “AZ” (Arizona) or “IA” (Iowa). QTL are colored by trait category. Confidence intervals for QTL are plotted as one-LOD interval. Genomic ranges for each LG are specified. For specific locations on the *G. hirsutum* genome sequence, LOD scores, and other details, see Table 3 and <u>Supplemental Table 2</u>.

**Table 2.** Subgenome location of linkage group based on linkage map and genomically mapped markers. The number of markers used to identify the chromosomes is listed. Start and end show the position in the corresponding *G. hirsutum* cv. TM-1 subgenome.

| Linkage group (AZ) | Linkage group (IA) | *G. hirsutum* | start | end | *G. arboreum* | *G. raimondii* |
| --- | --- | --- | --- | --- | --- | --- |
| AZ30 | IA24 | ChrA01 | 4,271,138 | 100,276,588 | Chr01/Chr02 |  |
| AZ25 |  | ChrA02 | 326,615 | 84,855,696 | Chr03 |  |
|  | IA11 | ChrA02 | 3,870,558 | 84,855,696 | Chr03 |  |
|  | IA12 | ChrA02 | 326,615 | 1,008,410 | Chr03 |  |
| AZ10 | IA07 | ChrA03 | 7,756,446 | 101,464,731 | Chr03 |  |
| AZ33 | IA32 | ChrA04 | 807,278 | 75,497,922 | Chr06 |  |
| AZ06 | IA16 | ChrA05 | 32,455,072 | 93,933,072 | Chr05 |  |
| AZ11 | IA34 | ChrA05 | 12,447,798 | 17,185,964 | Chr05 |  |
| AZ05 | IA06 | ChrA06 | 11,844,977 | 121,378,180 | Chr06 |  |
| AZ16 |  | ChrA07 | 1,830,647 | 89,848,877 | Chr06 |  |
| AZ17 |  | ChrA07 | 92,681,306 | 93,171,853 | Chr07 |  |
|  | IA22 | ChrA07 | 7,321,899 | 93,171,853 | Chr07 |  |
| AZ23 | IA19 | ChrA08 | 2,877,637 | 117,527,721 | Chr08 |  |
| AZ24 | IA05 | ChrA09 | 2,580,082 (15,659,999) | 79,333,397 (75,848,634) | Chr09 |  |
| AZ19 | IA15 | ChrA10 | 6,056,379 (6,566,496) | 106,114,506 | Chr10 |  |
| AZ08 | IA26 | ChrA11 | 1,912,510 | 4,371,131 | Chr11 |  |
| AZ15 |  | ChrA11 | 10,951,928 | 109,621,794 | Chr11 |  |
|  | IA17 | ChrA11 | 53,172,447 | 103,552,230 | Chr11 |  |
|  | IA18 | ChrA11 | 10,951,928 | 12,955,059 | Chr11 |  |
| AZ01 | IA02 | ChrA12 | 785,478 | 78,273,367 (72,842,063) | Chr12 |  |
| AZ03 | IA01 | ChrA12 | 77,411,923 (13,521,801) | 100,079,948 | Chr12 |  |
| AZ18 | IA08 | ChrA13 | 3,404,007 | 96,773,239 | Chr13 |  |
| AZ13 | IA10 | ChrD01 | 18,196,452 | 62,287,774 |  | Chr02 |
| AZ27 | IA33 | ChrD02 | 12,742,894 | 61,010,129 |  | Chr05 |
| AZ28 | IA36 | ChrD03 | 6,483,364 | 50,172,131 (48,393,682) |  | Chr03 |
| AZ14 | IA14 | ChrD04 | 3,602,330 | 56,438,319 |  | Chr12 |
| AZ12 |  | ChrD05 | 2,523,538 | 63,761,721 |  | Chr09 |
|  | IA27 | ChrD05 | 2,523,538 | 18,861,200 |  | Chr09 |
|  | IA28 | ChrD05 | 32,622,237 | 63,761,721 |  | Chr09 |
|  | IA29 | ChrD05 | 26,606,552 | 27,776,136 |  | Chr09 |
| AZ31 | IA31 | ChrD06 | 57,362,695 | 65,851,264 |  | Chr10 |
| AZ21 | IA20 | ChrD07 | 5,155,281 (18,304,091) | 48,192,327 |  | Chr01 |
| AZ22 | IA21 | ChrD07 | 55,033,970 | 55,696,530 |  | Chr01 |
| AZ09 | IA04 | ChrD08 | 2,309,559 (4,206,266) | 69,750,855 |  | Chr04 |
| AZ20 | IA23 | ChrD09 | 1,234,789 | 40,676,126 |  | Chr06 |
| AZ32 | IA30 | ChrD10 | 13,976,894 | 62,550,932 |  | Chr11 |
| AZ07 | IA25 | ChrD11 | 7,839,868 | 72,873,302 |  | Chr07 |
| AZ02 | IA03 | ChrD12 | 22,239,698 | 53,411,834 (51,612,631) |  | Chr08 |
| AZ04 | IA37 | ChrD12 | 61,838,133 | 101,355,435 |  | Chr08 |
| AZ26 | IA09 | ChrD13 | 8,757,166 | 58,413,467 |  | Chr13 |
| AZ29 | IA13 | ChrD13 | 62,947,661 |  |  | Chr13 |
| AZ34 | IA35 | ChrD13 | 852,543 | 1,182,162 |  | Chr05 |
| * | https://www.cottongen.org/species/Gossypium_hirsutum/jgi-AD1_genome_v1.1 |  |  |  |  |  |

**Table 3:** Top 50 QTL associated with domestication traits. For full list of QTL, see <u>Supplemental Table 2</u>.

| Category | Trait<sup>1</sup> | Chr<sup>2</sup> | QTL name<sup>3</sup> | Marker interval | Peak position (cM) | Peak position (Mb)<sup>4</sup> | LOD | A<sup>5</sup> | D<sup>6</sup> | \|d/a\|<sup>7</sup> | GA<sup>8</sup> | R<sup>2</sup>(%)<sup>9</sup> |
| --- | --- | --- | --- | --- | --- | --- | --- | --- | --- | --- | --- | --- |
| Fruiting habit | TN | A01 | qTN-AZ30-1 | c4_78149-EST1A_32413_01 | 22.20 | 65.15 | 8.68 | -2.28 | -0.97 | 0.42 | PD | 12.68 |
| Flower | PC | A05 | qPC-IA16-1 | c2_114307-c2_48932 | 36.07 | 32.46 | 8.13 | -0.11 | 0.11 | 1.02 | D | 13.82 |
| Fiber color | Ca | A06 | qCa-AZ5-1 | GS1A_19003p436q20-EST1A_111998 | 6.72 | 17.16 | 90.69 | -2.27 | 0.15 | 0.07 | A | 75.47 |
| Fiber color | Ca | A06 | qCa-IA6-1 | GS1A_14865p560q19-c4_48216 | 1.01 | 17.16 | 66.58 | 7.63 | -1.63 | 0.21 | PD | 75.40 |
| Fiber color | Cb | A06 | qCb-AZ5-1 | GS1A_19003p436q20-EST1A_111998 | 6.72 | 17.16 | 99.53 | -5.22 | 1.22 | 0.23 | PD | 79.89 |
| Fiber color | Cb | A06 | qCb-IA6-1 | GS1A_14865p560q19-c4_48216 | 1.01 | 17.16 | 55.90 | -2.23 | 0.45 | 0.20 | PD | 43.81 |
| Fiber color | CL | A06 | qCL-AZ5-1 | GS1A_19003p436q20-EST1A_111998 | 6.72 | 17.16 | 59.81 | 6.76 | -0.39 | 0.06 | A | 65.20 |
| Fiber length | L5n | A06 | qL5n-AZ5-1 | GS1A_19003p436q20-EST1A_116921 | 5.72 | 17.16 | 6.36 | 0.03 | 0.04 | 1.06 | D | 12.14 |
| Fiber length | Lw | A06 | qLw-AZ5-1 | GS1A_19003p436q20-EST1A_111998 | 5.72 | 17.16 | 6.23 | 0.03 | 0.02 | 0.68 | PD | 11.66 |
| Plant architecture | SP | A06 | qSP-AZ5-1 | GS1A_14865p560q19-c4_09782 | 17.84 | 96.62 | 72.39 | 1.48 | 0.15 | 0.10 | A | 71.49 |
| Plant architecture | SP | A06 | qSP-IA6-1 | GS1A_14865p560q19-EST1A_111998 | 11.14 | 100.61 | 43.49 | 1.20 | 0.02 | 0.01 | A | 48.49 |
| Fiber length | UQLw | A06 | qUQLw-AZ5-1 | GS1A_19003p436q20-EST1A_116921 | 6.72 | 17.16 | 5.54 | 0.03 | 0.03 | 1.03 | D | 12.13 |
| Fiber length | Lw | A07 | qLw-IA22-1 | c4_49169-cs_1083 | 69.75 | 72.79 | 5.43 | -0.03 | 0.00 | 0.03 | A | 10.79 |
| Other fiber qualities | MR | A07 | qMR-IA22-1 | GS1A_20202p545q14-c4_32659 | 67.17 | 72.79 | 4.13 | 0.66 | 1.45 | 2.18 | OD | 12.84 |
| Flower | PS | A07 | qPS-AZ16-1 | EST2_39330-c4_00014 | 26.01 | 21.29 | 62.23 | -0.38 | 0.51 | 1.33 | OD | 41.40 |
| Flower | PS | A07 | qPS-IA22-1 | c2_11322-GS1A_23155p1125q16 | 93.82 | 18.38 | 37.42 | -0.38 | 0.33 | 0.87 | D | 53.58 |
| Seed | SW | A07 | qSW-AZ16-1 | c4_32659-GS1A_23155p1125q16 | 18.01 | 80.66 | 8.69 | 0.24 | 0.00 | 0.01 | A | 12.87 |
| Fiber color | Ca | A08 | qCa-AZ23-1 | c4_21262-c4_44618 | 46.67 | 116.77 | 25.47 | -0.87 | 0.02 | 0.02 | A | 12.93 |
| Fiber color | CL | A08 | qCL-AZ23-1 | c4_21262-c4_44618 | 47.67 | 116.77 | 14.98 | 2.55 | 0.15 | 0.06 | A | 11.44 |
| Flower | CS | A12 | qCS-IA1-1 | c3_76188-GS2A_37259p664q5 | 2.91 | 78.27 | 8.20 | -0.26 | -0.22 | 0.85 | D | 25.99 |
| Fruiting habit | TN | A12 | qTN-IA2-1 | EST1A_00152-c4_13563 | 17.36 | 7.83 | 4.55 | -1.40 | 1.97 | 1.41 | OD | 14.07 |
| Fiber color | CL | D07 | qCL-IA20-1 | ck_75214-GS2A_20396p609q3 | 26.29 | 18.30 | 5.00 | -0.03 | 0.00 | 0.07 | A | 12.28 |
| Fiber length | L5n | D07 | qL5n-IA20-1 | ck_75214-GS2A_20396p609q3 | 28.29 | 18.30 | 4.49 | 2.65 | 1.65 | 0.62 | PD | 10.43 |
| Fiber length | LnCV | D07 | qLnCV-IA20-1 | ck_75214-GS2A_20396p609q3 | 28.29 | 18.30 | 4.49 | 2.65 | 1.65 | 0.62 | PD | 10.43 |
| Fruiting habit | PHTN | D07 | qPHTN-AZ21-1 | c4_46170-ck_75214 | 9.95 | 28.62 | 6.85 | -3.41 | -0.62 | 0.18 | A | 11.69 |
| Other fiber qualities | SFCn | D07 | qSFCn-IA20-1 | ck_75214-GS2A_20396p609q3 | 29.29 | 18.30 | 4.62 | 2.64 | 1.60 | 0.61 | PD | 10.77 |
| Fiber length | L25n | D01 | qL25n-AZ13-1 | EST1D_12_476-c4_21328 | 0.01 | 18.79* | 5.33 | -0.04 | 0.04 | 0.98 | D | 14.26 |
| Fiber length | L5n | D01 | qL5n-AZ13-1 | EST1D_12_476-c4_21328 | 7.27 | 18.20 | 8.39 | -0.05 | 0.00 | 0.05 | A | 14.27 |
| Fiber length | UQLw | D01 | qUQLw-AZ13-1 | EST1D_12_476-c4_21328 | 6.27 | 18.20 | 5.77 | -0.04 | 0.00 | 0.10 | A | 10.82 |
| Other fiber qualities | Fine | D09 | qFine-IA23-1 | c3_66591-c4_101926 | 53.97 | 12.45 | 4.13 | -1.50 | -2.56 | 1.71 | OD | 14.59 |
| Plant architecture | PHFB1 | D09 | qPHFB1-AZ20-1 | c3_66591-ck_77717 | 26.97 | 38.62 | 5.51 | 15.64 | -12.40 | 0.79 | PD | 10.49 |
| Fiber length | Lw | D11 | qLw-IA25-1 | c4_41050-c3_11416 | 55.76 | 20.78 | 5.28 | 0.04 | 0.00 | 0.08 | A | 11.10 |
| Fruiting habit | NF | D11 | qNF-AZ7-1 | c4_18678-c4_38231 | 101.27 | 10.42 | 7.53 | -1.09 | -0.76 | 0.70 | PD | 34.95 |
| Fruiting habit | TN | D11 | qTN-AZ7-1 | c4_18678-c4_38231 | 104.27 | 10.42 | 4.38 | -2.06 | -0.70 | 0.34 | PD | 11.56 |
| Fiber length | UQLw | D11 | qUQLw-IA25-1 | c4_41050-c3_11416 | 54.76 | 20.78 | 6.14 | 0.05 | -0.01 | 0.29 | PD | 13.61 |
| Flower | CS | D12 | qCS-IA37-1 | c4_41113-GS2A_7310p1793q4A | 0.01 | 61.84 | 36.03 | -0.43 | -0.55 | 1.28 | OD | 66.09 |
| Flower | CS | D05 | qCS-IA29-1 | EST1D_23510_01-GS2D_51331p736q38 | 3.01 | 27.78 | 30.54 | 0.41 | -0.54 | 1.31 | OD | 64.96 |
| Phenology | FBFF | D05 | qFBFF-IA27-1 | c4_06690-GS1D_23044p418q14 | 0.01 | 18.86 | 4.79 | -2.08 | -2.07 | 1.00 | D | 14.85 |
| Fruiting habit | TNFB | D05 | qTNFB-IA27-1 | c4_09461-c4_131801 | 61.46 | 9.09 | 4.33 | -1.32 | -1.03 | 0.79 | PD | 10.31 |
| Other fiber qualities | TrS | D05 | qTrS-AZ12-1 | EST2D_20572-GS1D_23044p418q14 | 92.71 | 18.86 | 5.11 | 19.84 | -7.19 | 0.36 | PD | 14.06 |
| Fiber length | UQLw | D05 | qUQLw-AZ12-1 | c2_04598-c4_131801 | 125.40 | 9.09 | 4.71 | -0.03 | 0.05 | 1.54 | OD | 17.51 |
| Plant architecture | SP | D06 | qSP-IA31-1 | EST1D_42236-EST1D_14_1412 | 45.29 | 55.33* | 14.64 | 0.61 | 0.20 | 0.33 | PD | 12.47 |
| Plant architecture | FB1 | D10 | qFB1-IA30-1 | GS1D_35377p470q22-ck_01124 | 6.01 | 62.55 | 4.05 | 0.13 | 1.26 | 9.64 | OD | 12.33 |
| Fiber length | L5n | D10 | qL5n-IA30-1 | ck_51389-c4_38839 | 26.55 | 13.98 | 5.15 | 2.34 | -2.69 | 1.15 | D | 11.51 |
| Fiber length | LnCV | D10 | qLnCV-IA30-1 | ck_51389-c4_38839 | 26.55 | 13.98 | 5.15 | 2.34 | -2.69 | 1.15 | D | 11.51 |
| Fiber length | Ln | D04 | qLn-AZ14-1 | EST1D_03860-c4_07376 | 80.59 | 3.60 | 4.40 | -0.03 | -0.01 | 0.33 | PD | 10.93 |
| Flower | PC | D04 | qPC-AZ14-1 | c4_02071-c2_50716 | 37.10 | 46.08* | 2.77 | -0.10 | 0.10 | 0.99 | D | 10.58 |
| Flower | PC | D04 | qPC-AZ14-2 | cs_12499-c4_07376 | 80.59 | 3.60 | 11.48 | -0.12 | 0.14 | 1.13 | D | 20.11 |
| Flower | PC | D04 | qPC-IA14-1 | EST1D_03860-c4_00820 | 10.61 | 39.70 | 7.73 | -0.10 | 0.13 | 1.28 | OD | 13.90 |
| Fiber length | L25n | D13 | qL25n-AZ26-1 | c4_18034-c2_58393 | 19.68 | 58.41 | 3.88 | 0.05 | 0.00 | 0.08 | A | 10.76 |

<sup>1</sup>Fiber color: Ca, mean a\*; Cb, mean b\*; CL, mean L\*; Fiber length: L25n, 2.5% Length by Number; L5n, 5% Length by Number; Ln, Mean Length by Number; LnCV, Coefficient of Variation of the Length by Number; Lw, Mean Length by Weight; UQLw, Upper Quantile Length by Weight; Flower: CS, Curly Style; PC, Pollen Color; PS, Petal Spot; Fruiting habit: NF, Total Number of Nodes to First Fruiting Branch; PHTN, Plant Height-to-Total Number of Nodes Ratio; TN, Total Number of Nodes; TNFB, Total Number of Non-Fruiting Branches; Other fiber qualities: Fine, Fineness; MR, Maturity Ratio; SFCn, Short Fiber Content by Number; TrS, Trash Size; Phenology: FBFF, Total Number of Fruiting Branches at First Flower; Plant architecture: FB1, Fruiting Branch Length for 1st Branch; PHFB1, Ratio of PH to FB1; SP, Average Stem Pubescence; Seed: SW, 50 Fuzzy Seed Weight.
<sup>2</sup>Chromosome designation. A and D represents the A- and D- subgenome, respectively.
<sup>3</sup>QTL name is provided as follows: the first two to four letters excluding “q” indicate the abbreviated trait name, following by linkage group (LG). The last letter indicates the population in which the QTL was detected; IA, Iowa; AZ, Arizona.
<sup>4</sup>Positions marked with an \* indicate estimates based on nearest genomically located markers.
<sup>5</sup>Additive (A) effect when substituting a TX2094 allele with an allele from Acala Maxxa at the QTL. The effect of the Acala Maxxa allele relative to the TX2094 allele at each QTL indicates the sign (positive or negative) of the allelic effect.
<sup>6</sup>Dominance (D) effect.
<sup>7</sup>|dominance effect/additive effect|
<sup>8</sup>Gene action. A, additive ( $|d/a| = 0-0.2$ ); PD, partial dominance ( $|d/a| = 0.21-0.8$ ); D, dominance ( $|d/a| = 0.81-1.2$ ); OD, overdominance ( $|d/a| > 1.2$ ).
<sup>9</sup>Percentage of phenotypic variance explained by each QTL.

### Candidate gene searches

Linkage groups were assigned to *G. hirsutum* chromosomes (Table 2) using molecular marker sequences as gmap (Wu and Watanabe 2005; Wu and Nacu 2010) queries against the published *G. hirsutum* cv TM-1 (CottonGen; Saski *et al.* 2017) genome (annotation gff version 1.1), using default values and permitting two possible paths (to accommodate homoeologs). A consensus of markers was used to identify the candidate chromosome for each linkage group, using the highest scoring path for each marker; however, when both paths were equally likely, both were used to derive the consensus. Candidate genes contained within the QTL confidence interval were identified by using the genomic coordinates of the first and last marker for each linkage group as a boundary, and subsequently intersecting the genomic boundaries of each linkage group with the genome annotation via bedtools 2 (Quinlan and Hall 2010). Orthogroups between the *G. hirsutum* genome used here and other published cotton genomes were generated via Orthofinder (Emms and Kelly 2015, 2019). Orthogroup results are not reported, but are provided for reference in Supplemental File 1. All scripts and parameters are available at https://github.com/Wendellab/QTL_TxMx.

Candidate genes were further screened for previously established expression differences in developing fibers (Bao, Hu et al, in revision), for putative transcription factors (CottonGen; Saski *et al.* 2017), and for non-silent SNPs between the parental accessions. For the latter, reads derived from *G. hirsutum* Acala Maxxa (SRA:SRR617482) and *G. hirsutum* TX2094 (SRA:SRR3560138-3560140) were mapped against the TM-1 genome (CottonGen; Saski *et al.* 2017) and SNPs were annotated using the Best Practices pipeline of GATK (Van der Auwera *et al.* 2013). The resulting vcf files were processed with vcftools (Danecek *et al.* 2011) and SnpSift (Cingolani *et al.* 2012a) to (1) only recover sites with differences between *G. hirsutum* Acala Maxxa and *G. hirsutum* TX2094, (2) remove sites with missing data, and (3) only recover SNPs where the wild *G. hirsutum* TX2094 shared the ancestral SNP with an outgroup species, *G. mustelinum* (SRA: SRR6334743). The resulting 3.6 million SNPs were annotated with SnpEff (Cingolani *et al.* 2012b) for the putative effects of each change, and SnpSift was again used to restrict the final vcf to only those SNPs where an effect was annotated. In addition, previously identified selective sweeps found in another *G. hirsutum* cv TM1 genome version (Fang *et al.* 2017a; Wang *et al.* 2017b) were placed on the *G. hirsutum* cv TM1 used here by comparing the genomes with MUMMER (Marçais *et al.* 2018) and intersecting coordinates with bedtools2 (Quinlan 2014). The final set of genes with annotated effects was further limited to only those regions under a QTL. These genes were additionally classified as to whether they also: (1) exhibit differential expression; (2) are putative TFs; or (3) belong to a curated list of potentially fiber-relevant cotton genes, based on existing literature (Fang 2018). Putative functional annotations were downloaded from CottonGen. The QTL peak was placed on the genome sequence by using the genomic QTL boundaries (determined above) to relate the number of cM to the amount of sequence in that same region (in base pairs). All program run information and relevant parameters are available at https://github.com/Wendellab/QTL_TxMx.

All data and scripts are available via GitHub (https://github.com/Wendellab/QTL_TxMx). Supplemental files are available at FigShare (<u>URL</u>) and on GitHub. All other data, e.g., genomes and downloaded sequences are listed in the methods. Seed from the mapping population is available from the GRIN National Genetic Resources Program.

## Results

### Phenotypic variation

Most traits investigated (Table 1) exhibited phenotypic variability between two parents, TX2094 and Acala Maxxa (<u>Supplemental Table 1</u>). In general, the phenotypes reflected the expected “domestication syndrome” in Acala Maxxa, as represented by its: (1) reduced plant height; (2) fewer total nodes; (3) fewer nodes to first fruiting branch; (4) better fruiting habit (e.g., longer fruiting branches); (5) early flowering; (6) greater production of flowers, bolls, and seeds; and (7) enhanced fiber quantity and quality (<u>Supplemental Table 1</u>). The F_2_ plants displayed a wide range of phenotypic variability in two greenhouse environments, Ames, IA, and Maricopa, AZ. The northern latitude of Iowa contributed to variability for traits reflective of a cooler, less-sunny environment compared to the F_2_ plants grown in Arizona. That is, plants grown in Iowa typically were taller, with shorter fruiting branch lengths and a greater number of nodes; however, these plants also exhibited a greater number of nodes to first fruiting branch, as well as a higher ratio of non-fruiting to fruiting branches. Interestingly, the Iowa subpopulation also exhibited both later flowering and more flowers during a 30-day interval. The flowers themselves exhibited greater distance between stigma and style, and produced more seeds per boll with an overall lighter seed weight (per boll), indicative of smaller seed size. Other flower and fiber traits exhibited continuous variation in all the F_2_ plants, from TX2094-like to Acala Maxxa-like phenotypes; however, the two subpopulations were often statistically distinguishable. For example, 50 Fuzzy Seed weight (g) was 3.96 and 4.13 in Iowa and Arizona, respectively, which is significantly different (*α* = 0.05). Observations such as these are unexpected under the null hypothesis that subpopulations should not be phenotypically distinct, and they likely reflect an interaction with the environment. Phenotypic measurements for parents and progeny are found in <u>Supplemental Table 1</u>.

### Linkage map construction

KASPar-based SNP genotyping was used to construct separate genetic linkage maps (total genetic length of 1704.03 cM for the Arizona subpopulation and 1989.46 cM for the Iowa subpopulation) from the *G. hirsutum* F_2_ subpopulations using JoinMap (Stam 1993). Of the 384 markers used for genotyping, 356 were successfully mapped to create 34 linkage groups for the Arizona population, and 336 were mapped to create 37 linkage groups for the Iowa population (Table 2). Among those 384 originally targeted markers, 84 markers were homoeolog-specific by design (see Byers et al. 2012). To determine whether the homoeologous genome of these markers was specific and accurately identified, linkage groups with multiple homoeolog-diagnostic SNPs were examined for genome consensus. Seventy (83%) of the 84 assays resided in linkage groups with at least one other homoeologous assay. The homoeologous genome assignment for these linkage groups was consistent with the genome sequence and the candidate gene/chromosome identification (see below). These linkage groups cover all 26 chromosomes in the *G. hirsutum* genome (Table 2).

### Identification of QTL and QTL clusters

A total of 120 QTL were detected from marker-trait analysis of the two subpopulations (Figure 2, <u>Supplemental Table 2</u>). The QTL detected from the subpopulations represented all phenotypic categories (53 QTL for 28 traits in the Iowa population; 67 QTL for 29 traits in the Arizona population). These QTL map to 22 and 24 linkage groups (20 and 21 chromosomes) in the Arizona and Iowa subpopulations, respectively; 59 QTL mapped to 12 chromosomes of A_T_ subgenome, while 61 QTL mapped to 12 chromosomes of D_T_ subgenome (<u>Supplemental Table 2</u>). In general, these *G. hirsutum* chromosomes carry a mean and median of 5 and 5.5 QTL respectively; however, three chromosomes (A02, A09 and A13) have only a single QTL each and two (A06, A07) include 10 QTL each (<u>Supplemental Table 2</u>). Combining QTL mapping results from two subpopulations, 11 QTL clusters were identified for 23 traits in eight trait categories (<u>Supplemental Table 2</u>). Seven QTL hotspots were identified on chromosomes A06 and A08 for fiber color, and chromosomes A6, A7, D01, D04 and D13 for fiber length (<u>Supplemental Table 2</u>). The top 50 QTL (R^2^ > 10%) are summarized in Table 3. A full listing of identified QTL, map, and genomic information, and other relevant information is included in <u>Supplemental Tables 2 and 3</u>, and is discussed in the context of phenotype (see below).

#### Connection of QTL to domestication

Of the 120 QTL identified across the two subpopulations, Acala Maxxa had additive allelic effects that were positive (‘increasing allele’) or negative (‘decreasing allele’), relative to Tx2094, for 56 and 64 QTL, respectively (<u>Supplemental Table 2</u>). With respect to trait, Acala Maxxa had more positive effect alleles for the 14 QTL (10 positive vs. 4 negative effect alleles) and 16 QTL (14 positive vs. 2 negative effect alleles) associated with traits in the plant architecture and seed categories. In contrast, Acala Maxxa had more QTL with negative allelic effects for traits in the fruiting habit (3 positive vs. 9 negative), flower (2 positive vs. 15 negative), and phenology (1 positive vs. 6 negative) categories. Interestingly, Acala Maxxa exhibited a more balanced number of positive and negative allelic effect estimates for the fiber length (16 positive vs. 17 negative), fiber color (5 positive vs, 8 negative), and other fiber qualities (5 positive vs. 3 negative). Collectively, these findings show that the QTL alleles contained within Acala Maxxa that associate with “domestication syndrome” attributes (e.g., greater production of seed, reduced stature, increased fiber length) may influence the phenotype in a manner not readily apparent (e.g., both positive and negative alleles associated with fiber length).

#### Candidate Gene identification

A total of 28,531 genes (<u>Supplemental Table 4</u>) are predicted within the genomic range of the 120 QTL (<u>Supplemental Table 2</u>), representing approximately 42% of the predicted gene models for the *G. hirsutum* cv. TM1 genome (Saski *et al.* 2017). The genomic regions occupied by QTL average approximately 83 Mbp in size (median = 76 Mbp), for a total genomic length of approximately 1,353 Mbp or 60% of the total sequenced genome length of 2,260 Mbp (<u>Supplemental Table 3</u>). For each phenotype (e.g., plant architecture, fiber color, etc), between 1,782-11,807 distinct genes were recovered. Candidate genes for each phenotype are discussed below.

We further screened the 28,531 candidate genes for (1) genes with non-silent mutations in the domesticated Acala Maxxa (using the outgroup polyploid species *G. mustelinum* to infer the ancestral state), to filter for possible functional differences at the protein level; (2) genes with expression differences between Acala Maxxa and TX2094, to filter for genes that have been up- or down-regulated under domestication; (3) transcription factors; or (4) known cotton fiber genes of interest (see methods for details) (<u>Supplemental Table 4</u>). In general, fewer genes were found within the QTL boundaries for the A subgenome (13,185 versus 15,346 in D_T_); while seemingly incongruent with the larger proportion of the A subgenome covered by QTL (approximately 847 Mbp in A_T_ versus 506 in D_T_), this likely reflects gene density differences due to the twofold difference in subgenome size (A ∼ 2D).

From the genome-wide total of 34,870 genes that have one or more SNP between TX2094 and Acala Maxxa, 87% (30,337 genes) are affected by at least one putatively non-silent mutation. Over half of these genes have SNPs that change the amino acid (19,195 genes), and slightly more than half have changes in the untranslated regions (UTR; 19,829) in an approximately 3:5 ratio favoring mutations in the 5′ UTR. These are slightly greater than the number of genes that have silent SNPs (39%; 13,579 genes). Only 2.6% of genes have a SNP that changes the start or stop (in an approximate 2:3 ratio, start:stop). Genome-wide, there exists no bias toward the A or D subgenome for any of the above categories. Of those 30,337 genes with non-silent TX2094 versus Acala Maxxa SNPs, 42% (12,744 genes) fall within a QTL in a ratio of approximately 0.8 A_T_:1 D_T_ (5,832 genes in A_T_ versus 6,912 in D_T_). This ratio is approximately equivalent to the overall representation of the genome under QTL, i.e., 0.9A_T_:1D_T_. Of the 12,744 genes with a non-silent SNP that occur under the QTL, 62% (7,925 genes) have predicted amino acid changes between TX2094 and Acala Maxxa (3,600 A_T_ genes and 4,325 D_T_) that could potentially be visible to selection (Table 4).

**Table 4:** Number of genes in any QTL, or for QTL related to a specific trait, that also exhibit additional differences between wild and domesticated cotton

|  |  |  |  |  |  |  |
| --- | --- | --- | --- | --- | --- | --- |
|  | Total | Genes with non-silent changes * | Genes with non-synonymous changes | differentially expressed ** | Transcription factors | Known cotton genes |
| All QTL | 28,531 | 12,744 | 1,617 | NA | 176 | 42 |
| Architecture | 5,646 | 2,602 | 490 | NA | 32 | 6 |
| Fiber Color | 1,782 | 764 | 3247 | 144 | 11 | 5 |
| Fiber Length | 11,807 | 5,254 | 1,230 | 865 | 80 | 16 |
| Other fiber qualities | 4203 | 1,963 | 2370 | 342 | 30 | 3 |
| Flower | 8,272 | 3816 | 1472 | NA | 50 | 14 |
| Fruiting Habit | 5,136 | 2335 | 813 | NA | 31 | 6 |
| Phenology | 2,661 | 1,297 | 2409 |  | 17 | 1 |
| Seed | 9,116 | 3,929 | 921 | NA | 54 | 15 |
| * includes start/stop adjustments and SNPs in UTR |  |  |  |  |  |  |
| ** DGE only applies to fiber-related traits |  |  |  |  |  |  |

To further explore the candidate genes under the QTL, we also quantified the number of genes under QTL that exhibit differential expression (DGE) during fiber development (Bao, Hu, et al. in press). Of the 5,168 genes differentially expressed between TX2094 and Acala Maxxa (in either 10 or 20 dpa fiber; adjusted *P*-value < 0.005), approximately 42% (2,148, genes) are located under one of the QTL (Table 4), over half of which were located under a fiber QTL (1,147). Between 7-8% of genes for each phenotypic group experienced DGE in the fiber stages surveyed (10 and 20 dpa). Interestingly, there appears to be little bias toward differential expression of genes under fiber-related QTL versus non-fiber QTL for these fiber-derived expression data. This may reflect a general overlap between fiber-relevant genes (e.g., cell wall, cytoskeletal genes, etc) and those involved in broad plant phenotypes, as well as the remarkable increase in gene coregulation during domestication (Hu *et al.* 2016). Therefore, while we note differences in DGE for possible candidate genes from any trait category, the relevance of this fiber-derived DGE to non-fiber traits is unclear. Differentially expressed genes that also contain nonsynonymous and/or UTR SNPs account for about half of the DGE-QTL genes (1,137 genes), 723 of which have predicted amino acid changes.

Finally, we also considered two categories of genes of possible interest under the QTL: transcription factors (TF) and previously identified fiber-relevant genes (see methods). The QTL regions contained 176 putative TF (CottonGen; Saski *et al.* 2017) (74A:102D), representing approximately 1% of the genes related to each trait. Of these 176 TF, 97 had putative amino acid changes. Only three transcription factors under QTL exhibited expression changes, i.e., Gohir.A04G012200 (qLw-IA32-1), Gohir.D05G036400 (qUQLw-AZ12-1 and qTNFB-IA27-1), and Gohir.D08G140800 (qLw-AZ9-1), which are mostly associated with fiber length (<u>Supplemental Table 2</u>). We also screened the genes underlying QTL for a compilation of 88 genes mined from the fiber biology literature (see methods). Of these, approximately half (42/88) were found under one or more QTL. Less than 1% of each phenotypic category was composed of genes derived from this list.

#### Plant architecture

Fourteen QTL were detected for 7 of 10 traits related to plant architecture on 10 chromosomes, 64% of which were from the Arizona population. Nearly half (6) of the fourteen QTL detected relate to stem pubescence, representing four distinct genomic locations and chromosomes; the remaining traits with QTL had only 1-2 QTL each. Particularly notable were the SP QTL located on chromosome A06 (linkage groups IA6 and AZ5), which explained 48.5 and 71.5% of the SP phenotypic variation, respectively. One QTL for plant height (PH) was detected in the D_T_-subgenome (D07; AZ21) in Arizona population, which explained 7.2% of the phenotypic variation (R^2^) and showed additivity. For PH, the TX2094 allele contributes to increasing height, although the two parental alleles work additively (Table 3; <u>Supplemental Table 2</u>).

Homology search of markers associated with these QTL identified 5,646 non-redundant genes in the QTL regions for plant architecture (<u>Supplemental Table 4</u>), with a mean of 433 genes per QTL. For plant height (PH), candidates include (Table 5), among others:a phototropic-responsive NPH3 family protein (Christie *et al.* 2018); a YUC8-like gene (Hentrich *et al.* 2013b); an auxin-responsive family protein (Gallavotti 2013); and tandem duplicates similar to putative far-red impaired responsive (FAR1) family proteins (Tang *et al.* 2013). Approximately 10% of the genes contained within the QTL exhibit differential expression between TX2094 and Maxxa, including a QUASIMODO-like homolog, which leads to a dwarf plant phenotype in Arabidopsis (Orfila *et al.* 2005). Fruiting branch-related traits exhibited 1-2 QTL for branch length (FB1, FB2) and *Plant Height-to-Fruiting Branch Length Ratio* (PHFB1, PHFB2). Interestingly, all QTL for FB1 and PHFB1 were found on D-derived chromosomes, whereas the QTL for FB2 and PHFB2 were found on A-derived chromosomes. Three phototropic-responsive NPH3-like genes are also found within these QTL (Table 5), which have demonstrated roles in *Arabidopsis* phototropism (Christie *et al.* 2018). Also contained within an FB2 QTL is an MKK7-like gene, which is implicated in plant architecture in *Arabidopsis* (*Wang and Li 2006*), while the single QTL for PHFB1 contains two tandem BIN2-like genes, which can affect plant height in *Arabidopsis* (Li 2005).

**Table 5:**
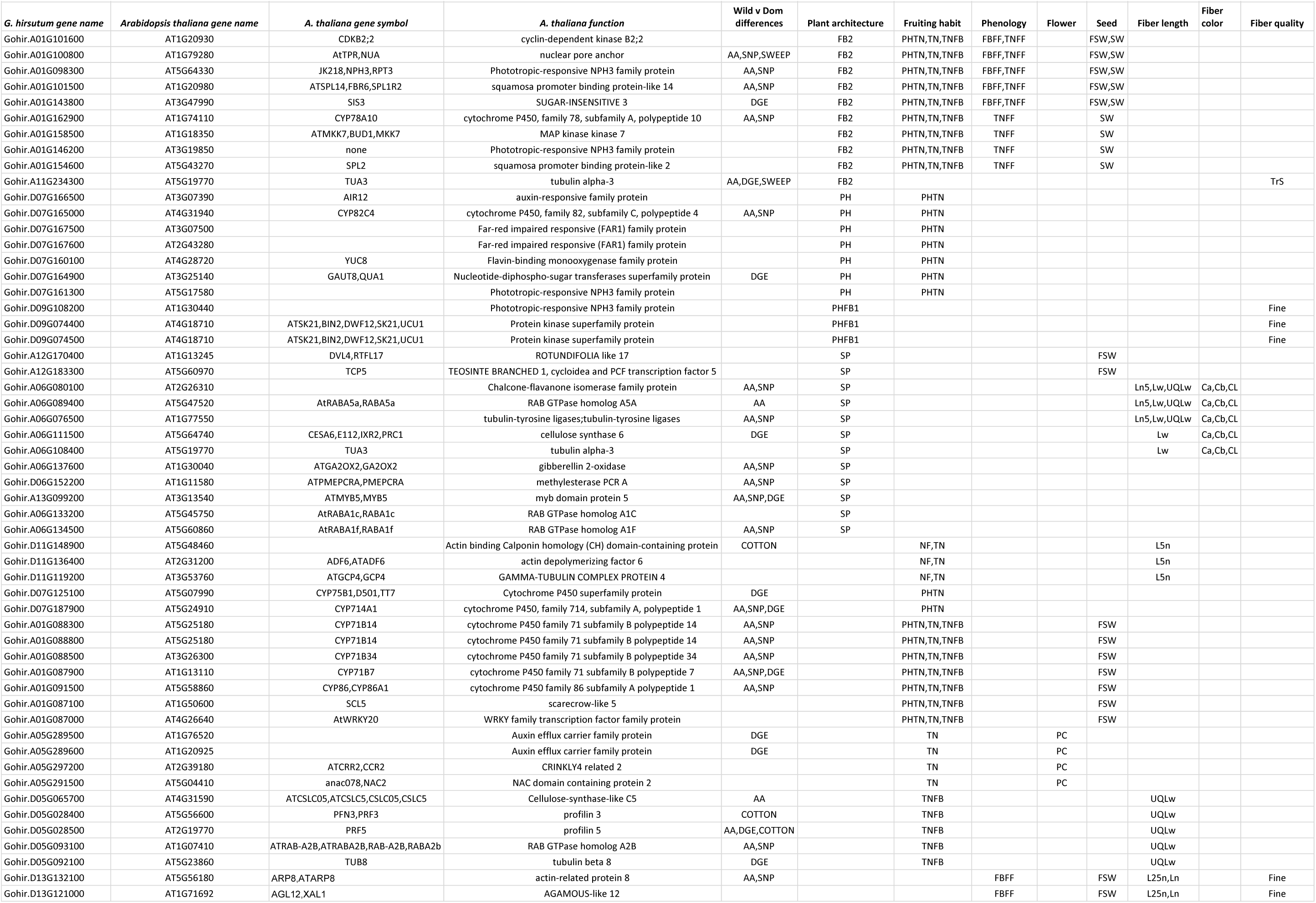
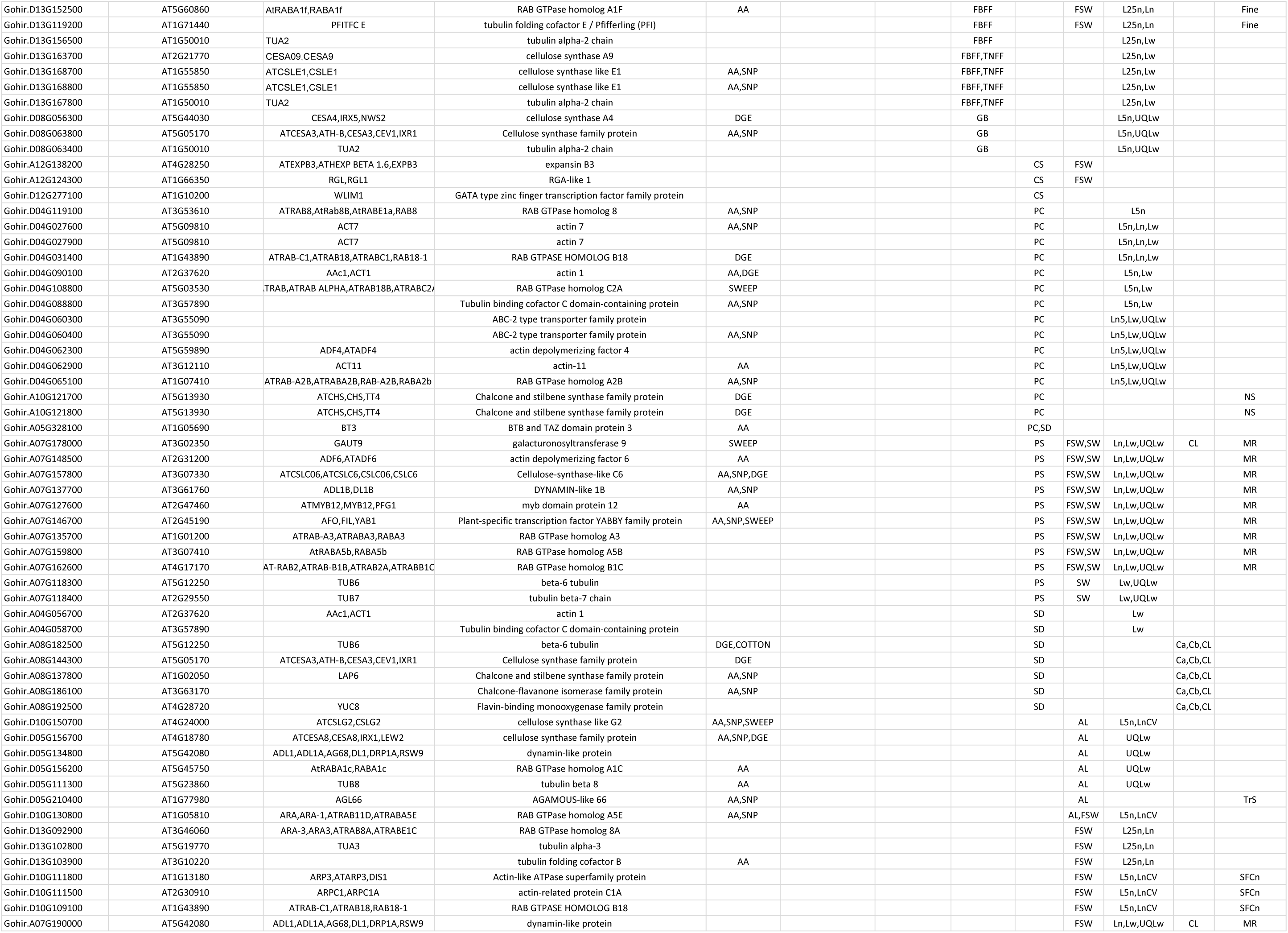
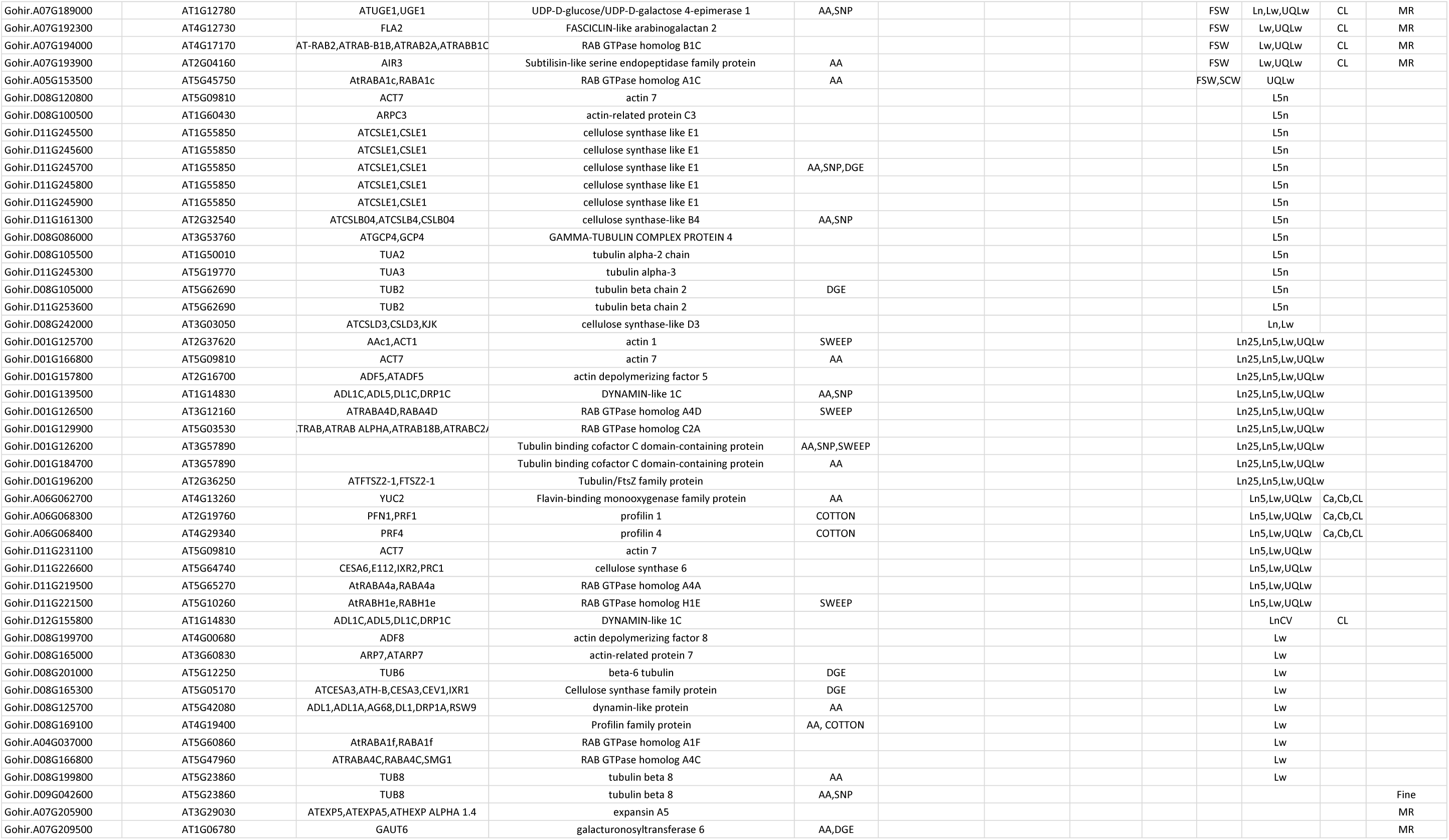
Possible candidates of interest. G. hirsutum gene name and closest Arabidopsis homolog are given (see methods for details). Candidates with amino acid (AA), non-silent SNP (SNP), gene expression (DGE) differences between wild and domesticated cotton are noted in column 5, as are known cotton genes with domestication effects (COTTON) or identified within regions of selective sweeps (SWEEP). Trait categories are listed in columns 6-13, and the traits with QTL that contain that gene are listed.

Stem pubescence had both the highest number of QTL and candidate genes, many of which have predicted functions in trichome and/or cell wall development, as well as amino acid changes between TX2094 and Acala Maxxa. One candidate is a predicted Myb 5-like gene (Table 5), which functions in trichome development in *Arabidopsis*. Two other candidates include two RAB GTPase-like genes, a gibberellin 2-oxidase-like gene, and a methylesterase-like gene, all of which have amino acid changes; genes involved in these processes are associated with cell wall metabolism or related pathways in *Arabidopsis* (Lycett 2008; Bischoff *et al.* 2010) and cotton (Xiao *et al.* 2019). Although somewhat further from the QTL peak, a cellulose synthase 6-like gene was found within the SP QTL, which is relevant to trichome development (Haigler *et al.* 2009; Betancur *et al.* 2010; Nixon *et al.* 2016)

#### Fruiting habit and Phenology

Nineteen QTL were detected for seven traits related to fruiting habit (4 traits) and phenology (3 traits; see Table 1), split evenly between subgenomes and scattered across 10 chromosomes. Five and three Fruiting Habit QTL were identified for Total Number of Nodes (TN) and Plant Height-to-Total Number of Nodes Ratio (PH_by_TN), respectively, in the Iowa and Arizona populations (<u>Supplemental Table 2</u>). Most QTL for PH_by_TN showed additivity, whereas only one exhibited additivity for TN; the remaining four QTL exhibited partial- or over-dominance. Three QTL were detected for Total Number of Non-Fruiting Branches (TNFB) dispersed across three chromosomes (2 A_T_ and 1 D_T_) and occurring in both subpopulations (2 Iowa, 1 Arizona), whereas a single QTL was found for Total Number of Nodes to First Fruiting Branch (NF) in the Arizona subpopulation, which was found on chromosome D11 and explained 35% of the variation for the trait.

Two phenology QTL were identified for Total Number of Nodes at First Flower (TNFF) in the Iowa population only. The two QTL for TNFF were either partial or over-dominance and explained ∼7% of the phenotypic variation each, whereas the three QTL for FBFF were either dominant, overdominant, or additive, explaining between 7.9-14.9% of the variation. Interestingly, while the final Phenology trait, Total Number of Green Bolls Retained after 30 days + 4 week interval (GB) exhibited two QTL (Arizona subpopulation only), one from each subgenome, the chromosomes were not homoeologous (i.e, were not homologous in the diploid progenitors).

Homology searches of QTL-associated markers recovered 5,136 non-redundant genes in the QTL intervals controlling fruiting habit and 2,661 genes in the intervals controlling phenology. Although many of the same chromosomes were implicated in both trait categories, only 714 genes are shared between the two. Nearly half of the genes recovered for both traits exhibited SNPs with potential effects (e.g., amino acid changes) between TX2094 and Acala Maxxa (45% and 49% for Fruiting Habit and Phenology, respectively); however, few genes exhibited differential expression (8% in each; <u>Supplemental Table 4</u>). Putative candidates for PH_by_TN include two genes similar to *Arabidopsis* WRKY and GRAS transcription factors (Table 5) and at least nine cytochrome P450-like genes, which are part of a relatively large superfamily of genes with diverse metabolic roles (Mizutani and Ohta 2010; Mizutani 2012); most of these cytochrome P450-like genes (6) have predicted amino acid changes between TX2094 and Acala Maxxa.Total number of nodes (TN) QTL candidate genes include two differentially expressed auxin efflux carrier family proteins; a differentially expressed SIS3-like homolog; and a CCR-related gene (Table 5). Homologs of SIS3 are involved in the growth response to high concentrations of exogenous sugars (Huang *et al.* 2010)members of the CCR gene family may be involved in lignin biosynthesis during development (Lauvergeat *et al.* 2001). Several genes are found associated with the TN QTL in regions that overlap the TNFB QTL, including a homolog of SPL2, which is involved in shoot maturation and the transition to flowering (Shikata *et al.* 2009); a nuclear pore anchor, whose *Arabidopsis* homolog affects flowering time regulation and other developmental processes (Xu *et al.* 2007); and two adjacent genes, a squamosa promoter binding protein-like and a cyclin-dependent kinase B2;2-like gene,, both of which are involved in plant growth and development (Andersen *et al.* 2008; Jorgensen and Preston 2014). For the single QTL involved in NF, no obvious candidate genes were noted; however, 46% of the 660 genes in the QTL regions were affected by non-conservative SNPs (see methods), including 29% with amino acid changes. Interestingly, many Fruiting habit QTL candidates overlap those found in Plant architecture (Table 5), which may reflect an overlap in developmental programmes.

While three traits representing the Phenology trait category each recovered QTL (i.e., FBFF, GB, and TNFF), the QTL for FBFF and TNFF largely overlapped. Most QTL regions encompassed by TNFF were also found for FBFF, except for part of chromosome A01, where the FBFF QTL is more narrowly predicted than in TNFF. This region of chromosome A01 also has many overlapping QTL for Fruiting habit and other Phenology traits (i.e., PHTN, TN, TNFB), which may indicate that it is a notable region for plant growth and development. The other QTL for FBFF were located solely on the D_T_ chromosomes, and includes an AGAMOUS-like gene (Table 5), which could act responsively to plant hormones and have function in regulating fruit formation in cotton (de Moura *et al.* 2017). Interestingly, the QTL for FBFF on chromosome D13 overlaps with QTL for Fiber Length and therefore contains some fiber-relevant genes (Table 5), including a tubulin-related gene. Similarly, one of the two QTL for GB entirely overlaps with 1-2 Fiber length QTL on chromosome D08, while the other QTL completely overlaps with the Plant Architecture QTL PHFB2 (see above). These overlapping QTL regions may also reflect overlap in developmental programmes between fiber development, plant architecture and growth, and fruit retention.

#### Flower

Seventeen QTL were identified for four floral traits, which individually explain 4.6-66.1% of the phenotypic variation and most of which exhibited varying degrees of dominance. Four QTL were detected for Average Stigma Distance (SD), two from each population, on four different chromosomes (A04, A05, A08 and D11). Four QTL were also identified for Curly Style (CS) from the Iowa population only, with the curly allele typically originating from TX2094. Seven QTL were detected for Pollen Color (PC) on two A and two D chromosomes (A05, A10, D04, and D05); presence of TX2094 alleles generated more yellow pollen (<u>Supplemental Table S2</u>). Finally, two QTL were detected for the presence of a petal spot (PS; chromosome A07), a TX2094-derived trait.

Candidate gene searches revealed 8,272 genes in the QTL intervals for floral traits. The QTL for curly style exhibited several genes related to cell wall formation and/or organization, which may be involved in conferring the curly phenotype (Table 5). These include an RGA-like gene that may play a role in regulating organ development (Wang *et al.* 2009); an expansin B3-like gene which may be involved in cell wall expansion mediation (Shcherban *et al.* 1995; Lee *et al.* 2001); and a WLIM1-like transcription factor whose *Arabidopsis* homolog regulates cytoskeletal organization via interaction with actin filaments (Papuga *et al.* 2010). Likewise, several notable genes were detected for pollen color. Two of these are arrayed in tandem and are putative ABC-2 type transporter-like genes; this gene family participates in pollen wall synthesis, as observed in *Arabidopsis* (Yadav *et al.* 2014). A second tandem array of two putative homologs of chalcone synthase was also found for PC, with both members exhibiting differential expression between Acala Maxxa and TX2094 (albeit measured in fiber only). An additional PC-related gene is an NAC-like gene with a possible role in regulating flavonoid biosynthesis (Morishita *et al.* 2009). Similarly, the single notable gene within the QTL for PS is a myb domain protein whose *Arabidopsis* homolog is involved in flavonoid biosynthesis (Wang *et al.* 2016b). The QTL for average stigma distance includes a single gene of interest, a transcription factor which plays a role in male and female gametophyte development (Robert *et al.* 2009).

#### Seed

Sixteen QTL were identified representing five of the seven seed-related traits (<u>Supplemental Table 2</u>), which individually explain 5.6-12.87% of the variance per trait. The trait 50 Fuzzy Seed Weight (FSW) had the most QTL (7), distributed over 6 chromosomes. The remaining traits had 1-3 associated QTL, most having a positive effect allele from the domesticated Acala Maxxa parent. Most seed QTL reside on A_T_ subgenome chromosomes (10 out of 16, including 5 of the QTL for FSW).

QTL for Seed-related traits contain 9,116 candidate genes. For the fuzzy seed weight QTL regions, these include a UDP-D-glucose/-galactose 4-epimerase and several FASCICLIN-like arabinogalactans (FLA), including a FLA2-like gene (Table 5). Both of these exhibit up-regulation in domesticated (versus wild) cottons (Yoo and Wendel 2014) and have *Arabidopsis* homologs that function in cell wall biosynthesis. Also included in theQTL region is a Pfifferling (PFI)-like homolog, which functions in seed (embryo) development in *Arabidopsis* (Steinborn *et al.* 2002), and an expansion (EXPA5)-like homolog, which may act to mediate cell wall expansion (Shcherban *et al.* 1995; Lee *et al.* 2001). Notably, these genes all belong to the FSW QTL, which overlaps in these regions with QTL for fiber traits. An additional two candidate genes within the FSW QTL have possible roles in fruit formation: a DVL-homolog that may confer phenotypic changes in fruit and inflorescence (Wen *et al.* 2004), and an AGAMOUS 12-like gene whose family has a suggested role in cotton fruit formation (de Moura *et al.* 2017). The only other notable candidate gene within the Seed QTL is another AGAMOUS-like gene, which was found within the QTL for AL.

#### Fiber length

Fiber-related characteristics were among the obvious phenotypic targets during domestication of cotton. Not surprisingly, therefore, 54 QTL were detected for fiber-related traits (i.e., length, color, and measures of quality), of which 33 (61%) were for fiber length (<u>Supplemental Table 2</u>). As observed in some other populations, a majority of these (76% or 25 QTL) were located in the subgenome (D_T_) derived from the parental diploid that has short, unspinnable fiber. These QTL were dispersed over 9 of the 13 D_T_ chromosomes and 4 of the 13 A_T_-derived chromosomes, individually explaining from 7.2 to 17.5% of the phenotypic variation. Despite having far fewer QTL, the A_T_-subgenome exhibited QTL for four of the seven length traits evaluated (<u>Supplemental Table 2</u>). Only 4 of the A_T_-subgenome QTL explained more than 10% of the variation (versus 12 D_T_ QTL) and only one was in the top 5 fiber-length related QTL, explaining at most 12.1% of the trait variation. Conversely, nearly half of the QTL found on D_T_-subgenome chromosomes (<u>Supplemental Table 2</u>) individually explain over 10% of the phenotypic variation (R^2^) for their categories (12 out of 25 D_T_ QTL).

Candidate gene searches for fiber length QTL revealed several possibilities (Table 5), including 19 cellulose synthase-like genes, most of which (17) are found on the D_T_ chromosomes and five of which clustered on chromosome D11. The middle gene in this cluster, Gohir.D11G245700, exhibited both amino acid changes and differential gene expression between wild and domesticated *G. hirsutum*, supporting a possible role in fiber domestication. Differential expression was also found for four other cellulose synthase-like genes, including both genes found on the A_T_ chromosomes. Because many of the fiber QTL overlap, nearly half (8) of the cellulose synthase genes were associated with multiple Fiber length QTL (mean=1.5 QTL). Interestingly, an additional cellulose synthase-like gene (Gohir.A08G144300) was also differentially expressed between wild and domesticated cotton; however, this gene was not contained within any fiber length QTL, but was rather found associated with multiple fiber color QTL and one for Average Stigma Distance (<u>Supplemental Table 4</u>). Similarly, several genes typically associated with flavonoid production (e.g., chalcone-flavanone isomerase) were found within the fiber length QTL rather than the QTL for fiber color where they would be expected to influence the brown coloration found in wild fibers.

As expected, many additional candidate genes involved in cytoskeleton/cell wall formation or trichome development were found, including several genes with known associations with fiber development (Table 5). Twenty-five tubulin related genes were found associated with fiber length QTL, including eight beta tubulin-like genes. Beta tubulin genes are relevant to cell wall development because they orient the cellulose microfibrils (Spokevicius *et al.* 2007), a major component of secondary cell walls. Three of the beta tubulin-like genes exhibit differential expression between wild and domesticated cotton fiber, and each is associated with a different QTL trait (Table 5). Eighteen actin-related genes were also found within the fiber QTL, including one with a known role in fiber elongation and secondary wall synthesis (Gohir.D11G148900; (Zhang *et al.* 2017)); however, no differential expression or SNPs with predicted functional consequences were detected between wild and domesticated cotton for this gene. Five profilin homologs were associated with fiber length; profilin expression has previously been associated with fiber domestication (Bao *et al.* 2011). Six dynamin(DL1)-like proteins were also associated with Fiber length, along with 22 RAB GTPase-like genes (Table 5). In *Arabidopsis,* these genes influence cell wall composition (both) and cellular expansion (DL1) (Collings *et al.* 2008). Notably, the DL1-like candidate and one RAB GTPase-like candidate exhibits differential expression between wild and domesticated cotton fiber. Finally, a YABBY1 transcription factor-like gene was associated with fiber length whose *Arabidopsis* homoeolog is exclusively expressed in trichomes (Schliep *et al.* 2010). This candidate gene also exhibits an amino acid change between wild and domesticated cotton.

#### Fiber color

Fiber color is conferred by the accumulation of flavonoids in mature fibers (Hua *et al.* 2007; Xiao *et al.* 2007, 2014; Li *et al.* 2012a; Feng *et al.* 2013; Tuttle *et al.* 2015). Thirteen QTL were detected for the three fiber color traits evaluated: mean *L\** (bright/dark), mean *a\** (green/red), and mean *b\** (blue/yellow). Many of these on chromosomes A06 and A08 overlapped between populations and traits, and therefore aggregate into two distinct QTL hotspots. The QTL on chromosome A06 were typically of major effect, individually explaining from 43.8 to 79.9% of the phenotypic variation, whereas those on chromosome A08 typically explained less than 10% of the variation (from 5.1 to 12.9%; mean 8.8 %). Two flavin-binding monooxygenase family (YUCCA)-like proteins were found within the color QTL detected here, one each on chromosomes A06 and A08 (Table 5). *Arabidopsis* homologs of the YUCCA family function in the production of auxin (Hentrich *et al.* 2013a, 2013b), a key regulator of plant development that may also be involved in the regulation of flavonol synthesis (Lewis *et al.* 2011). Likewise, a chalcone-flavanone isomerase family-like protein was found within the color QTL on both A06 and A08, which also functions in flavonoid biosynthesis in *Arabidopsis* (Jiang *et al.* 2015). Chromosome A08 has an additional flavonol-related candidate gene, i.e., a chalcone and stilbene synthase family protein. Interestingly, while chromosomes A06 and A08 have loci with predicted relevance to fiber color, the QTL on chromosomes A07, D07, and D12 do not exhibit any notable candidates; however, the color QTL for chromosomes A07 and D12 do overlap QTL for fiber length and fiber quality in which there exist several genes that may influence fiber morphology (Table 5). These include the previously mentioned dynamin-like gene, a gene similar to FASCICLIN-like arabinogalactan that has been implicated in fiber domestication (Yoo and Wendel 2014) and cell wall biosynthesis (MacMillan *et al.* 2010), and a TUB6-like gene. Whether the overlap of these QTL is coincidence or suggests an overlap in the genetic networks conferring different fiber traits is unknown and will require future research on the fiber development network.

#### Other fiber qualities

While a total of 14 “other” measures of fiber quality were evaluated (Table 1), only five traits produced QTL (8 QTL), namely, Fineness, Maturity Ratio, Nep Size, Short Fiber Content by Number, and Trash Size. Each trait was associated with 1-2 QTL each for a total of 8 QTL located on as many chromosomes. Several candidates affecting cell wall composition and synthesis were found within these two regions (Table 5). These include two tubulin-like genes, Gohir.A11G234300 and Gohir.D09G042600, which exhibit differential expression and amino acid changes, respectively. An actin-like ATPase found in this region is similar to the *Arabidopsis* ARP3 gene, which controls trichome shape (Mathur *et al.* 2003). The region also includes a subtilisin protease-like candidate; subtilisin proteases have been associated with cell wall composition in *Arabidopsis thaliana*, specifically the mucilage content of cell walls (Rautengarten *et al.* 2008). Two additional candidates are galacturonosyltransferase (GAUT)-like genes (Table 5), whose *Arabidopsis thaliana* homologs influence cell wall composition by controlling pectin biosynthesis (Caffall 2008; Caffall *et al.* 2009; Atmodjo *et al.* 2011).

### Comparison of putative QTL between subpopulations, between subgenomes, and among chromosomes

The F_2_ seed derived from a single cross between *G. hirsutum* accessions TX2094 and Acala Maxxa were planted in two different greenhouse environments, in Maricopa, AZ and Ames, IA (see methods). The 120 total QTL detected were nearly evenly divided between the two subpopulations, with Arizona recovering slightly more QTL (67 QTL, or 56%) than Iowa. While the number of QTL recovered in each subpopulation was similar, only 22 QTL were declared as coincident QTL between the two locations, and eight of them shared peak markers. Likewise, while both populations detected QTL on a similar number of chromosomes (20 and 21 in Arizona and Iowa, respectively), approximately 30% of chromosomes (7) had QTL from only one population. On average, the QTL detected in Iowa had a slightly more narrow range (<u>Supplemental Table 2</u>), both overall (13.2 versus 19.1 cM, or 14 versus 39 Mb) and when only considering QTL regions with the same peak marker (18.6 versus 20.7 cM, or 5 versus 30 Mb). Slight and opposing subgenome biases were found for the chromosomes recovered from each subpopulation, with Iowa recovering QTL on 11 A_T_ and 10 D_T_ chromosomes, whereas Arizona recovered QTL on 9 A_T_ and 11 D_T_ chromosomes.

The QTL peaks shared between the Iowa and Arizona subpopulations were exclusively associated with fiber color (2 peak markers, 4 QTL regions; <u>Supplemental Table 2</u>), with the remaining seven coincident regions influencing fiber length (1 shared QTL region), flower (3 shared QTL regions), seed (1 shared QTL regions), and plant architecture (2 shared QTL regions). Eight of the 11 coincident QTL regions were located on A_T_-derived chromosomes, with chromosome A06 represented most frequently (3 shared QTL regions; Figure 2). Three of the 8 trait categories surveyed had no shared QTL regions, i.e., Fiber Quality, Fruiting Habit, and Phenology; this is possibly due in part to these being the categories with the fewest QTL reported (<u>Supplemental Table 2</u>).

The distribution and total length of the 120 QTL was nearly equivalent between the two polyploid subgenomes (59A:61D); however, when QTL redundancy between subpopulations is considered, this proportion becomes slightly D-biased (51A:58D). This may be due to the bias toward A_T_ chromosomes in shared QTL and a slight overrepresentation of D_T_-derived QTL in the Arizona population (32A:35D). Both the mean and median length of A_T_ derived QTL are larger than for D_T_ derived QTL (36.5 versus 16 Mb, respectively, for mean, and 31 versus 8 Mb for median), which is likely a consequence of the larger genome size (twofold) inherited from the A diploid parent. Slightly more than half of the categories (i.e., fiber color, flower, fruiting habit, and seed) had more A_T_ QTL, with fiber color exhibiting the largest bias (85% A_T_-derived QTL). Fiber length exhibited the next greatest bias, albeit for the opposite subgenome; i.e., approximately 76% (25) of fiber length QTL are D_T_-derived. In fact, approximately half of the total D_T_-derived QTL are associated with fiber length (∼41% overall). Interestingly, because the fiber quality category also contained more D_T_-derived QTL (3A:5D), these two fiber categories together accounting for nearly half of the QTL from D_T_ subgenome chromosomes and over 73% of the QTL for these categories. This observation is congruent with some previous research that has suggested D-genome recruitment during fiber domestication.

## Discussion

### QTL lability and the complex genetic architecture of cotton domestication phenotypes

The molecular underpinnings of the domesticated cotton fiber phenotype are of substantial interest from both evolutionary and economic standpoints. Because a cotton “fiber” is a highly exaggerated single-celled structure, it provides a unique model for the evolutionary and developmental transformations that are possible in a single cell. Economically, cotton fibers are central to a multi-billion dollar and globally vital industry, one that has a vested interest in manipulating the genetics of domesticated fiber. Consequently, myriad studies have attempted to reveal the key players in fiber development. The results of these experiments and analyses have been diverse and often in conflict, underscoring the complex nature of cotton fiber biology and also the diverse suite of populations that have variously been employed. Comparison between the present research and previously generated QTL suffers from this same complexity. Many of the phenotypic traits evaluated here have been evaluated in other crosses and under different conditions, as summarized in the Cotton QTL Database v. 2.3 (Said *et al.* 2015a) and CottonGen (Yu *et al.* 2014). As noted by others, QTL results of an individual study (such as the one presented here) are frequently incongruent with QTL results from other crosses grown under different conditions (Rong *et al.* 2007; Lacape *et al.* 2010; Said *et al.* 2015b, 2015a). This observation is clear from our results alone, where less than half of the QTL were shared across two similar environments. When extended to previous QTL results, even our most robust QTL (*i.e.*, fiber color, chromosome A06) exhibit more complicated inheritance; *i.e.*, the Cotton QTL Database lists 62 QTL for fiber color spread across 21 of the 26 cotton chromosomes whereas we detect a single chromosome of major effect and only 4 of lesser effect for both environments. A notable difference between ours and previous studies, however, is that ours was designed to capture the array of changes that characterize the transformation of the truly wild form of *G. hirsutum* into the modern elite cultivars that presently comprise the modern annualized crop plant. This cross should capture the major differences between wild and domesticated forms of *G. hirsutum*, whereas previous research has focused on differences between either (1) elite lines of the independently domesticated species *G. hirsutum* and *G. barbadense* (i.e., Pima cotton), or (2) between *G. hirsutum* landraces and/or elite cultivars, which reflect differences in improvement rather than those accompanying initial domestication.

Notwithstanding these substantive differences among studies, both the results presented here and earlier indicate that the genetic architecture underlying fiber morphology and development (among other domestication phenotypes) is complex and is responsive to environmental conditions. Consequently, uncovering QTL represent an important yet insufficient step in disentangling the genetic underpinnings of fiber development and cotton domestication. The complex interactions among genes important to understanding the QTL recovered remain to be elucidated, but many important enabling tools for such analyses have been developed. For example, gene coexpression network analyses can reveal modules of interconnected genes involved in key traits, as shown for cottonseed (Hu *et al.* 2016) and fiber (Gallagher et al. in prep), using the comparative context of wild versus domesticated *G. hirsutum*. In these examples, domestication appears to have increased the coordinated expression among genes and gene modules relevant to domesticated phenotypes. Research on *cis*/*trans* regulatory differences between wild and domesticated *G. hirsutum* (Bao, Hu, et al. in press) indicates that changes in both *cis* and *trans* regulation have occurred during domestication, which are significantly enriched with fiber QTL genes reported here. Notably, regulatory variations are frequently associated with environmental responsiveness (Cubillos *et al.* 2014; Lovell *et al.* 2016; Waters *et al.* 2017) and therefore may underlie the environmental variability of QTL as reported.

### Multiple sources of information can narrow candidate gene identification

A primary goal of QTL analyses is to uncover the genomic basis of phenotypic differences. In many cases, QTL regions encompass a large region of the genome, and hence contain many genes. Here, each individual QTL recovered between 14 and 1,678 genes (mean = 531), resulting in 1,782 - 11,807 possible candidate genes for each phenotype (<u>Supplemental Table 2</u>). In the present analysis, we narrow the candidate genes to focus on those genes with secondary evidence, i.e., DGE, amino acid changes, transcription factors, and/or those with relevant functions in related species. The genes mentioned here as candidates, while not exhaustive, represent possible causative sources for the their respective phenotypes. The strength of these candidates, however, is limited by the information available. For the fiber QTL, we were able to leverage existing expression information for the accessions used in the QTL mapping cross, which provides additional evidence supporting individual genes as candidates. A caveat, however, is that since the expression sampling was completed for an independent project and QTL are often environmentally labile, genes exhibiting differential expression (or lack thereof) in the dataset used here may not represent the expression patterns that would be observed in the individuals used in the initial QTL cross and grown under the conditions of the QTL subpopulations. Furthermore, differential expression data were only available for two timepoints during fiber development, albeit key timepoints (Haigler *et al.* 2012). Future QTL research may be improved by integrating multiple data types from the outset, including expression from tissues relevant to the phenotypes evaluated for each parent grown in each environment; however, the results of the present were improved (for the fiber phenotype) by considering the data available.

### Implications for domestication and future prospects

Domestication is a complex process involving a multiplicity of traits and the coordinated alteration of gene expression for numerous genes, for all but the simplest of traits (Olsen and Wendel 2013a, 2013b; Meyer and Purugganan 2013; Kantar *et al.* 2017; Purugganan 2019). With respect to cotton, a large number of QTL analyses have been conducted, specifically focused on economically valuable fiber characteristics, with some interest in other agronomically important phenotypes. These analyses have used either different species (Jiang *et al.* 1998; Paterson *et al.* 2003; Mei *et al.* 2004; Lacape *et al.* 2005, 2010; Chee *et al.* 2005a, 2005b; Draye *et al.* 2005; Rong *et al.* 2007; Said *et al.* 2015b, 2015a; Wang *et al.* 2016a, 2017a, 2017c) or different cultivated lines of the same species (Ulloa *et al.* 2005; Zhang *et al.* 2005; Shen *et al.* 2006; Qin *et al.* 2008; Lin *et al.* 2009; Li *et al.* 2012b, 2013; Tang *et al.* 2015; Tan *et al.* 2015, 2018; Wang *et al.* 2015; Shang *et al.* 2015, 2016; Jamshed *et al.* 2016) to provide perspectives on the genetic control of various traits. While each contributes to our multi-dimensional understanding of the controls on phenotypes, (1) it is not immediately clear that interspecies QTL are useful in cotton breeding programs (Lin *et al.* 2009; Shang *et al.* 2015; Jamshed *et al.* 2016), and (2) inter-cultivar or inter-line crosses provide a limited perspective on the underlying genetic architecture leading to modern elite lines. The present QTL analysis was designed specifically to reveal the genetic architecture underlying the morphological transformation from wild to domesticated upland cotton, *G. hirsutum*. Like many of existing QTL analyses in cotton, our cross, while having allelic replication only in two environments, also demonstrates that the genomic differences that underlie many wild vs. cultivated characteristics are environmentally variable. Only about 18% of the QTL were shared across the two subpopulations. This variability is likely due to pleiotropic and environmentally labile regulatory factors and genetic interactions (Wittkopp *et al.* 2004; Coolon *et al.* 2014; Chen *et al.* 2015; Metzger *et al.* 2016; Rhoné *et al.* 2017; Signor and Nuzhdin 2018) playing a role in divergence between wild and domesticated species. This complexity is also increased by the allopolyploid nature of cotton, whose subgenomes evolved in isolation for 5-10 million years but now are reunited in a common nucleus, where they have coexisted for 1-2 million years. It is notable that, congruent with other QTL analyses, we find important fiber related QTL on the subgenome derived from the parent with the much shorter, inferior fiber (D genome). The involvement of the D-genome in the evolution of transgressive fiber phenotypes has been noted in multiple analyses, including for QTL (Jiang *et al.* 1998; Lacape *et al.* 2005; Han *et al.* 2006; Rong *et al.* 2007; Qin *et al.* 2008; Said *et al.* 2015b), expression (Hovav *et al.* 2008a; Yoo and Wendel 2014; Zhang *et al.* 2015; Fang *et al.* 2017b), and in selective genomic sweeps (Fang *et al.* 2017a, 2017c; Song *et al.* 2019), yet the underlying genetic basis for this phenomenon remains unclear. Further work using advanced populations in which individual QTL have been isolated in isogenic backgrounds, combined with a multi-omics or systems biology perspective, is one promising approach for developing a fuller understanding of cotton biology as well as the domestication process.

## Acknowledgements

The authors would like to thank Gwen Coyle, Mark Arick, Kristen Cox, Joel Gilley, Virginia Moreno, Anna Tuchin, and Kara Grupp for experimental assistance. The authors acknowledge computational support and assistance from the Iowa State University ResearchIT Unit (http://researchit.las.iastate.edu/). Funding was provided by the National Science Foundation Plant Genome Program, the USDA-ARS, and by Cotton Incorporated.

## References

Ai, X., Y. Liang, J. Wang, J. Zheng, Z. Gong et al., 2017 Genetic diversity and structure of elite cotton germplasm (Gossypium hirsutum L.) using genome-wide SNP data. Genetica 145: 409–416.

Al-Ghazi, Y., S. Bourot, T. Arioli, E. S. Dennis, and D. J. Llewellyn, 2009 Transcript profiling during fiber development identifies pathways in secondary metabolism and cell wall structure that may contribute to cotton fiber quality. Plant Cell Physiol. 50: 1364–1381.

Andersen, S. U., S. Buechel, Z. Zhao, K. Ljung, O. Novák et al., 2008 Requirement of B2-type cyclin-dependent kinases for meristem integrity in Arabidopsis thaliana. Plant Cell 20: 88–100.

Argiriou, A., A. Kalivas, G. Michailidis, and A. Tsaftaris, 2012 Characterization of PROFILIN genes from allotetraploid (Gossypium hirsutum) cotton and its diploid progenitors and expression analysis in cotton genotypes differing in fiber characteristics. Mol. Biol. Rep. 39: 3523–3532.

Atmodjo, M. A., Y. Sakuragi, X. Zhu, A. J. Burrell, S. S. Mohanty et al., 2011 Galacturonosyltransferase (GAUT)1 and GAUT7 are the core of a plant cell wall pectin biosynthetic homogalacturonan:galacturonosyltransferase complex. Proc. Natl. Acad. Sci. U. S. A. 108: 20225– 20230.

Bao, Y., G. Hu, L. E. Flagel, A. Salmon, M. Bezanilla et al., 2011 Parallel up-regulation of the profilin gene family following independent domestication of diploid and allopolyploid cotton (Gossypium). Proc. Natl. Acad. Sci. U. S. A. 108: 21152–21157.

Betancur, L., B. Singh, R. A. Rapp, J. F. Wendel, M. D. Marks et al., 2010 Phylogenetically distinct cellulose synthase genes support secondary wall thickening in arabidopsis shoot trichomes and cotton fiber. J. Integr. Plant Biol. 52: 205–220.

Bischoff, V., S. Nita, L. Neumetzler, D. Schindelasch, A. Urbain et al., 2010 TRICHOME BIREFRINGENCE and its homolog AT5G01360 encode plant-specific DUF231 proteins required for cellulose biosynthesis in Arabidopsis. Plant Physiol. 153: 590–602.

Brubaker, C. L., and J. F. Wendel, 1994 Reevaluating the Origin of Domesticated Cotton (Gossypium hirsutum; Malvaceae) Using Nuclear Restriction Fragment Length Polymorphisms (RFLPs). Am. J. Bot. 81: 1309–1326.

Byers, R. L., D. B. Harker, S. M. Yourstone, P. J. Maughan, and J. A. Udall, 2012 Development and mapping of SNP assays in allotetraploid cotton. Theor. Appl. Genet. 124: 1201–1214.

Caffall, K. H., 2008 Expression and characterization of galacturonosyltransferase-6 (GAUT6) of the galacturonosyltranserferase-1 (GAUT1)-related gene family of Arabidopsis thaliana.

Caffall, K. H., S. Pattathil, S. E. Phillips, M. G. Hahn, and D. Mohnen, 2009 Arabidopsis thaliana T-DNA mutants implicate GAUT genes in the biosynthesis of pectin and xylan in cell walls and seed testa. Mol. Plant 2: 1000–1014.

Cai, C., G. Zhu, T. Zhang, and W. Guo, 2017 High-density 80 K SNP array is a powerful tool for genotyping G. hirsutum accessions and genome analysis. BMC Genomics 18: 654.

Chandnani, R., Z. Zhang, J. D. Patel, J. Adhikari, S. Khanal et al., 2017 Comparative genetic variation of fiber quality traits in reciprocal advanced backcross populations. Euphytica 213: 241.

Chaudhary, B., R. Hovav, L. Flagel, R. Mittler, and J. F. Wendel, 2009 Parallel expression evolution of oxidative stress-related genes in fiber from wild and domesticated diploid and polyploid cotton (Gossypium). BMC Genomics 10: 378.

Chee, P., X. Draye, C.-X. Jiang, L. Decanini, T. A. Delmonte et al., 2005a Molecular dissection of interspecific variation between Gossypium hirsutum and Gossypium barbadense (cotton) by a backcross-self approach: I. Fiber elongation. Theor. Appl. Genet. 111: 757–763.

Chee, P. W., X. Draye, C.-X. Jiang, L. Decanini, T. A. Delmonte et al., 2005b Molecular dissection of phenotypic variation between Gossypium hirsutum and Gossypium barbadense (cotton) by a backcross-self approach: III. Fiber length. Theor. Appl. Genet. 111: 772–781.

Chen, J., V. Nolte, and C. Schlötterer, 2015 Temperature stress mediates decanalization and dominance of gene expression in Drosophila melanogaster. PLoS Genet. 11: e1004883.

Christie, J. M., N. Suetsugu, S. Sullivan, and M. Wada, 2018 Shining Light on the Function of NPH3/RPT2-Like Proteins in Phototropin Signaling. Plant Physiol. 176: 1015–1024.

Churchill, G. A., and R. W. Doerge, 1994 Empirical threshold values for quantitative trait mapping. Genetics 138: 963–971.

Cingolani, P., V. M. Patel, M. Coon, T. Nguyen, S. J. Land et al., 2012a Using Drosophila melanogaster as a Model for Genotoxic Chemical Mutational Studies with a New Program, SnpSift. Front. Genet. 3: 35.

Cingolani, P., A. Platts, L. L. Wang, M. Coon, T. Nguyen et al., 2012b A program for annotating and predicting the effects of single nucleotide polymorphisms, SnpEff: SNPs in the genome of Drosophila melanogaster strain w1118; iso-2; iso-3. Fly 6: 80–92.

Collings, D. A., L. K. Gebbie, P. A. Howles, U. A. Hurley, R. J. Birch et al., 2008 Arabidopsis dynamin-like protein DRP1A: a null mutant with widespread defects in endocytosis, cellulose synthesis, cytokinesis, and cell expansion. J. Exp. Bot. 59: 361–376.

Coolon, J. D., C. J. McManus, K. R. Stevenson, B. R. Graveley, and P. J. Wittkopp, 2014 Tempo and mode of regulatory evolution in Drosophila. Genome Res. 24: 797–808.

Coppens d’Eeckenbrugge, G., and J.-M. Lacape, 2014 Distribution and differentiation of wild, feral, and cultivated populations of perennial upland cotton (Gossypium hirsutum L.) in Mesoamerica and the Caribbean. PLoS One 9: e107458.

CottonGen Download TM-1 (G. hirsutum, AD1) Genome Data.

Cubillos, F. A., O. Stegle, C. Grondin, M. Canut, S. Tisné et al., 2014 Extensive cis-regulatory variation robust to environmental perturbation in Arabidopsis. Plant Cell 26: 4298–4310.

Danecek, P., A. Auton, G. Abecasis, C. A. Albers, E. Banks et al., 2011 The variant call format and VCFtools. Bioinformatics 27: 2156–2158.

Draye, X., P. Chee, C.-X. Jiang, L. Decanini, T. A. Delmonte et al., 2005 Molecular dissection of interspecific variation between Gossypium hirsutum and G. barbadense (cotton) by a backcross-self approach: II. Fiber fineness. Theor. Appl. Genet. 111: 764–771.

Emms, D. M., and S. Kelly, 2019 OrthoFinder: phylogenetic orthology inference for comparative genomics. bioRxiv 466201.

Emms, D. M., and S. Kelly, 2015 OrthoFinder: solving fundamental biases in whole genome comparisons dramatically improves orthogroup inference accuracy. Genome Biol. 16: 157.

Fang, D. D., 2018 Cotton Fiber: Physics, Chemistry and Biology. Springer.

Fang, L., H. Gong, Y. Hu, C. Liu, B. Zhou et al., 2017a Genomic insights into divergence and dual domestication of cultivated allotetraploid cottons. Genome Biol. 18: 33.

Fang, L., X. Guan, and T. Zhang, 2017b Asymmetric evolution and domestication in allotetraploid cotton (Gossypium hirsutum L.). The Crop Journal 5: 159–165.

Fang, L., Q. Wang, Y. Hu, Y. Jia, J. Chen et al., 2017c Genomic analyses in cotton identify signatures of selection and loci associated with fiber quality and yield traits. Nat. Genet. 49: 1089–1098.

Feng, H., X. Tian, Y. Liu, Y. Li, X. Zhang et al., 2013 Analysis of flavonoids and the flavonoid structural genes in brown fiber of upland cotton. PLoS One 8: e58820.

Fluidigm, 2011 SNP Genotyping:.

Fryxell, P. A., 1976 A Nomenclator of Gossypium: The Botanical Name of Cotton: United States Department of Agriculture, Economic Research Service Technical Bulletins 158582.

Fryxell, P. A., 1968 A Redefinition of the Tribe Gossypieae. Bot. Gaz. 129: 296–308.

Fryxell, P. A., 1992 A revised taxonomic interpretation of Gossypium L (Malvaceae). Rheeda 2: 108– 165.

Fryxell, P. A., 1979 Natural History of the Cotton Tribe. Texas A&M University Press.

Gallagher, J. P., C. E. Grover, K. Rex, M. Moran, and J. F. Wendel, 2017 A New Species of Cotton from Wake Atoll, Gossypium stephensii (Malvaceae). Syst. Bot. 42: 115–123.

Gallavotti, A., 2013 The role of auxin in shaping shoot architecture. J. Exp. Bot. 64: 2593–2608.

Gou, J.-Y., L.-J. Wang, S.-P. Chen, W.-L. Hu, and X.-Y. Chen, 2007 Gene expression and metabolite profiles of cotton fiber during cell elongation and secondary cell wall synthesis. Cell Res. 17: 422– 434.

Haigler, C. H., L. Betancur, M. R. Stiff, and J. R. Tuttle, 2012 Cotton fiber: a powerful single-cell model for cell wall and cellulose research. Front. Plant Sci. 3: 104.

Haigler, C. H., B. Singh, G. Wang, and D. Zhang, 2009 Genomics of Cotton Fiber Secondary Wall Deposition and Cellulose Biogenesis, pp. 385–417 in Genetics and Genomics of Cotton, edited by A. H. Paterson. Springer US, New York, NY.

Han, Z., C. Wang, X. Song, W. Guo, J. Gou et al., 2006 Characteristics, development and mapping of Gossypium hirsutum derived EST-SSRs in allotetraploid cotton. Theor. Appl. Genet. 112: 430–439.

Hentrich, M., C. Böttcher, P. Düchting, Y. Cheng, Y. Zhao et al., 2013a The jasmonic acid signaling pathway is linked to auxin homeostasis through the modulation of YUCCA8 and YUCCA9 gene expression. Plant J. 74: 626–637.

Hentrich, M., B. Sánchez-Parra, M.-M. Pérez Alonso, V. Carrasco Loba, L. Carrillo et al., 2013b YUCCA8 and YUCCA9 overexpression reveals a link between auxin signaling and lignification through the induction of ethylene biosynthesis. Plant Signal. Behav. 8: e26363.

Hinze, L. L., A. M. Hulse-Kemp, I. W. Wilson, Q.-H. Zhu, D. J. Llewellyn et al., 2017 Diversity analysis of cotton (Gossypium hirsutum L.) germplasm using the CottonSNP63K Array. BMC Plant Biol. 17: 37.

Hovav, R., B. Chaudhary, J. A. Udall, L. Flagel, and J. F. Wendel, 2008a Parallel domestication, convergent evolution and duplicated gene recruitment in allopolyploid cotton. Genetics 179: 1725– 1733.

Hovav, R., J. A. Udall, B. Chaudhary, R. Rapp, L. Flagel et al., 2008b Partitioned expression of duplicated genes during development and evolution of a single cell in a polyploid plant. Proc. Natl. Acad. Sci. U. S. A. 105: 6191–6195.

Hovav, R., J. A. Udall, E. Hovav, R. Rapp, L. Flagel et al., 2008c A majority of cotton genes are expressed in single-celled fiber. Planta 227: 319–329.

Huang, Y., C. Y. Li, D. L. Pattison, W. M. Gray, S. Park et al., 2010 SUGAR-INSENSITIVE3, a RING E3 ligase, is a new player in plant sugar response. Plant Physiol. 152: 1889–1900.

Hua, S., X. Wang, S. Yuan, M. Shao, X. Zhao et al., 2007 Characterization of Pigmentation and Cellulose Synthesis in Colored Cotton Fibers. Crop Sci. 47: 1540–1546.

Hu, Y., J. Chen, L. Fang, Z. Zhang, W. Ma et al., 2019 Gossypium barbadense and Gossypium hirsutum genomes provide insights into the origin and evolution of allotetraploid cotton. Nat. Genet. 51: 739– 748.

Hu, G., R. Hovav, C. E. Grover, A. Faigenboim-Doron, N. Kadmon et al., 2016 Evolutionary Conservation and Divergence of Gene Coexpression Networks in Gossypium (Cotton) Seeds. Genome Biol. Evol. 8: 3765–3783.

Jamshed, M., F. Jia, J. Gong, K. K. Palanga, Y. Shi et al., 2016 Identification of stable quantitative trait loci (QTLs) for fiber quality traits across multiple environments in Gossypium hirsutum recombinant inbred line population. BMC Genomics 17: 197.

Jiang, C., R. J. Wright, K. M. El-Zik, and A. H. Paterson, 1998 Polyploid formation created unique avenues for response to selection in Gossypium (cotton). Proc. Natl. Acad. Sci. U. S. A. 95: 4419– 4424.

Jiang, W., Q. Yin, R. Wu, G. Zheng, J. Liu et al., 2015 Role of a chalcone isomerase-like protein in flavonoid biosynthesis in Arabidopsis thaliana. J. Exp. Bot. 66: 7165–7179.

Jorgensen, S. A., and J. C. Preston, 2014 Differential SPL gene expression patterns reveal candidate genes underlying flowering time and architectural differences in Mimulus and Arabidopsis. Mol. Phylogenet. Evol. 73: 129–139.

Kantar, M. B., A. R. Nashoba, J. E. Anderson, B. K. Blackman, and L. H. Rieseberg, 2017 The Genetics and Genomics of Plant Domestication. Bioscience 67: 971–982.

Kaur, B., P. Tyagi, and V. Kuraparthy, 2017 Genetic Diversity and Population Structure in the Landrace Accessions of Gossypium hirsutum. Crop Sci. 57: 2457–2470.

Kim, H. J., B. A. Triplett, H.-B. Zhang, M.-K. Lee, D. J. Hinchliffe et al., 2012 Cloning and characterization of homeologous cellulose synthase catalytic subunit 2 genes from allotetraploid cotton (Gossypium hirsutum L.). Gene 494: 181–189.

Kutner, M., C. Nachtsheim, J. Neter, and W. Li, 2004 Applied Linear Statistical Models. McGraw-Hill/Irwin.

Lacape, J.-M., D. Llewellyn, J. Jacobs, T. Arioli, D. Becker et al., 2010 Meta-analysis of cotton fiber quality QTLs across diverse environments in a Gossypium hirsutum x G. barbadense RIL population. BMC Plant Biol. 10: 132.

Lacape, J.-M., T.-B. Nguyen, B. Courtois, J.-L. Belot, M. Giband et al., 2005 QTL Analysis of Cotton Fiber Quality Using Multiple Gossypium hirsutum Gossypium barbadense Backcross Generations. Published in Crop Sci. 45: 123–140.

Lauvergeat, V., C. Lacomme, E. Lacombe, E. Lasserre, D. Roby et al., 2001 Two cinnamoyl-CoA reductase (CCR) genes from Arabidopsis thaliana are differentially expressed during development and in response to infection with pathogenic bacteria. Phytochemistry 57: 1187–1195.

Lee, J. A., 1968 Genetical Studies concerning the Distribution of Trichomes on the Leaves of GOSSYPIUM HIRSUTUM L. Genetics 60: 567–575.

Lee, Y., D. Choi, and H. Kende, 2001 Expansins: ever-expanding numbers and functions. Curr. Opin. Plant Biol. 4: 527–532.

Lewis, D. R., M. V. Ramirez, N. D. Miller, P. Vallabhaneni, W. K. Ray et al., 2011 Auxin and ethylene induce flavonol accumulation through distinct transcriptional networks. Plant Physiol. 156: 144–164.

Li, J., 2005 Brassinosteroid signaling: from receptor kinases to transcription factors. Curr. Opin. Plant Biol. 8: 526–531.

Li, T., H. Fan, Z. Li, J. Wei, Y. Lin et al., 2012a The accumulation of pigment in fiber related to proanthocyanidins synthesis for brown cotton. Acta Physiol. Plant 34: 813–818.

Lin, Z., Y. Zhang, X. Zhang, and X. Guo, 2009 A high-density integrative linkage map for Gossypium hirsutum. Euphytica 166: 35–45.

Liu, B., and J. F. Wendel, 2002 Intersimple sequence repeat (ISSR) polymorphisms as a genetic marker system in cotton: TECHNICAL NOTE. Mol. Ecol. Notes 1: 205–208.

Li, C., C. Wang, N. Dong, X. Wang, H. Zhao et al., 2012b QTL detection for node of first fruiting branch and its height in upland cotton (Gossypium hirsutum L.). Euphytica 188: 441–451.

Li, C., X. Wang, N. Dong, H. Zhao, Z. Xia et al., 2013 QTL analysis for early-maturing traits in cotton using two upland cotton (*Gossypium hirsutum* L.) crosses. Breed. Sci. 63: 154–163.

Lovell, J. T., S. Schwartz, D. B. Lowry, E. V. Shakirov, J. E. Bonnette et al., 2016 Drought responsive gene expression regulatory divergence between upland and lowland ecotypes of a perennial C4 grass. Genome Res. 26: 510–518.

Lycett, G., 2008 The role of Rab GTPases in cell wall metabolism. J. Exp. Bot. 59: 4061–4074.

MacMillan, C. P., S. D. Mansfield, Z. H. Stachurski, R. Evans, and S. G. Southerton, 2010 Fasciclin-like arabinogalactan proteins: specialization for stem biomechanics and cell wall architecture in Arabidopsis and Eucalyptus: FLAs specialized for stem biomechanics and cell walls. Plant J. 62: 689–703.

Ma, Z., S. He, X. Wang, J. Sun, Y. Zhang et al., 2018 Resequencing a core collection of upland cotton identifies genomic variation and loci influencing fiber quality and yield. Nat. Genet. 50: 803–813.

Mangin, B., B. Goffinet, and A. Rebaï, 1994 Constructing confidence intervals for QTL location. Genetics 138: 1301–1308.

Marçais, G., A. L. Delcher, A. M. Phillippy, R. Coston, S. L. Salzberg et al., 2018 MUMmer4: A fast and versatile genome alignment system. PLoS Comput. Biol. 14: e1005944.

Mathur, J., N. Mathur, V. Kirik, B. Kernebeck, B. P. Srinivas et al., 2003 Arabidopsis CROOKED encodes for the smallest subunit of the ARP2/3 complex and controls cell shape by region specific fine F-actin formation. Development 130: 3137–3146.

McCarty, J. C., D. D. Deng, J. N. Jenkins, and L. Geng, 2018 Genetic diversity of day-neutral converted landrace Gossypium hirsutum L. accessions. Euphytica 214: 173.

McCouch, S. R., Y. G. Cho, P. E. Yano, M. Blinstrub, H. Morishima et al., 1997 Report on QTL nomenclature. Rice Genet. Newsl. 14: 11–13.

Mei, M., N. H. Syed, W. Gao, P. M. Thaxton, C. W. Smith et al., 2004 Genetic mapping and QTL analysis of fiber-related traits in cotton (Gossypium). Theor. Appl. Genet. 108: 280–291.

Metzger, B. P. H., F. Duveau, D. C. Yuan, S. Tryban, B. Yang et al., 2016 Contrasting Frequencies and Effects of cis- and trans-Regulatory Mutations Affecting Gene Expression. Mol. Biol. Evol. 33: 1131–1146.

Meyer, R. S., and M. D. Purugganan, 2013 Evolution of crop species: genetics of domestication and diversification. Nat. Rev. Genet. 14: 840–852.

Mizutani, M., 2012 Impacts of diversification of cytochrome P450 on plant metabolism. Biol. Pharm. Bull. 35: 824–832.

Mizutani, M., and D. Ohta, 2010 Diversification of P450 genes during land plant evolution. Annu. Rev. Plant Biol. 61: 291–315.

Morishita, T., Y. Kojima, T. Maruta, A. Nishizawa-Yokoi, Y. Yabuta et al., 2009 Arabidopsis NAC transcription factor, ANAC078, regulates flavonoid biosynthesis under high-light. Plant Cell Physiol. 50: 2210–2222.

de Moura, S. M., S. Artico, C. Lima, S. M. Nardeli, A. Berbel et al., 2017 Functional characterization of AGAMOUS-subfamily members from cotton during reproductive development and in response to plant hormones. Plant Reprod. 30: 19–39.

Nigam, D., P. Kavita, R. K. Tripathi, A. Ranjan, R. Goel et al., 2014 Transcriptome dynamics during fibre development in contrasting genotypes of Gossypium hirsutum L. Plant Biotechnol. J. 12: 204– 218.

Nixon, B. T., K. Mansouri, A. Singh, J. Du, J. K. Davis et al., 2016 Comparative Structural and Computational Analysis Supports Eighteen Cellulose Synthases in the Plant Cellulose Synthesis Complex. Sci. Rep. 6: 28696.

Olsen, K. M., and J. F. Wendel, 2013a A bountiful harvest: genomic insights into crop domestication phenotypes. Annu. Rev. Plant Biol. 64: 47–70.

Olsen, K. M., and J. F. Wendel, 2013b Crop plants as models for understanding plant adaptation and diversification. Front. Plant Sci. 4: 290.

Orfila, C., S. O. Sørensen, J. Harholt, N. Geshi, H. Crombie et al., 2005 QUASIMODO1 is expressed in vascular tissue of Arabidopsis thaliana inflorescence stems, and affects homogalacturonan and xylan biosynthesis. Planta 222: 613–622.

Papuga, J., C. Hoffmann, M. Dieterle, D. Moes, F. Moreau et al., 2010 Arabidopsis LIM proteins: a family of actin bundlers with distinct expression patterns and modes of regulation. Plant Cell 22: 3034–3052.

Paterson, A. H., Y. Saranga, M. Menz, C.-X. Jiang, and R. Wright, 2003 QTL analysis of genotype × environment interactions affecting cotton fiber quality. Theor. Appl. Genet. 106: 384–396.

Purugganan, M. D., 2019 Evolutionary Insights into the Nature of Plant Domestication. Curr. Biol. 29: R705–R714.

Qin, H., W. Guo, Y.-M. Zhang, and T. Zhang, 2008 QTL mapping of yield and fiber traits based on a four-way cross population in Gossypium hirsutum L. Theor. Appl. Genet. 117: 883–894.

Quinlan, A. R., 2014 BEDTools: the Swiss-army tool for genome feature analysis. Curr. Protoc. Bioinformatics 47: 11–12.

Quinlan, A. R., and I. M. Hall, 2010 BEDTools: a flexible suite of utilities for comparing genomic features. Bioinformatics 26: 841–842.

Rapp, R. A., C. H. Haigler, L. Flagel, R. H. Hovav, J. A. Udall et al., 2010 Gene expression in developing fibres of Upland cotton (Gossypium hirsutum L.) was massively altered by domestication. BMC Biol. 8: 139.

Rautengarten, C., B. Usadel, L. Neumetzler, J. Hartmann, D. Büssis et al., 2008 A subtilisin-like serine protease essential for mucilage release from Arabidopsis seed coats. Plant J. 54: 466–480.

Reddy, U. K., P. Nimmakayala, V. L. Abburi, C. V. C. M. Reddy, T. Saminathan et al., 2017 Genome-wide divergence, haplotype distribution and population demographic histories for Gossypium hirsutum and Gossypium barbadense as revealed by genome-anchored SNPs. Sci. Rep. 7: 41285.

Rhoné, B., C. Mariac, M. Couderc, C. Berthouly-Salazar, I. S. Ousseini et al., 2017 No Excess of Cis-Regulatory Variation Associated with Intraspecific Selection in Wild Pearl Millet (Cenchrus americanus). Genome Biol. Evol. 9: 388–397.

Robert, H. S., A. Quint, D. Brand, A. Vivian-Smith, and R. Offringa, 2009 BTB and TAZ domain scaffold proteins perform a crucial function in Arabidopsis development. Plant J. 58: 109–121.

Rong, J., F. A. Feltus, V. N. Waghmare, G. J. Pierce, P. W. Chee et al., 2007 Meta-analysis of polyploid cotton QTL shows unequal contributions of subgenomes to a complex network of genes and gene clusters implicated in lint fiber development. Genetics 176: 2577–2588.

Said, J. I., J. A. Knapka, M. Song, and J. Zhang, 2015a Cotton QTLdb: a cotton QTL database for QTL analysis, visualization, and comparison between Gossypium hirsutum and G. hirsutum × G. barbadense populations. Mol. Genet. Genomics 290: 1615–1625.

Said, J. I., Z. Lin, X. Zhang, M. Song, and J. Zhang, 2013 A comprehensive meta QTL analysis for fiber quality, yield, yield related and morphological traits, drought tolerance, and disease resistance in tetraploid cotton. BMC Genomics 14: 776.

Said, J. I., M. Song, H. Wang, Z. Lin, X. Zhang et al., 2015b A comparative meta-analysis of QTL between intraspecific Gossypium hirsutum and interspecific G. hirsutum × G. barbadense populations. Mol. Genet. Genomics 290: 1003–1025.

Saski, C. A., B. E. Scheffler, A. M. Hulse-Kemp, B. Liu, Q. Song et al., 2017 Sub genome anchored physical frameworks of the allotetraploid Upland cotton (Gossypium hirsutum L.) genome, and an approach toward reference-grade assemblies of polyploids. Sci. Rep. 7: 15274.

Schliep, M., B. Ebert, U. Simon-Rosin, D. Zoeller, and J. Fisahn, 2010 Quantitative expression analysis of selected transcription factors in pavement, basal and trichome cells of mature leaves from Arabidopsis thaliana. Protoplasma 241: 29–36.

Shang, L., Q. Liang, Y. Wang, X. Wang, K. Wang et al., 2015 Identification of stable QTLs controlling fiber traits properties in multi-environment using recombinant inbred lines in Upland cotton (Gossypium hirsutum L.). Euphytica 205: 877–888.

Shang, L., Y. Wang, X. Wang, F. Liu, A. Abduweli et al., 2016 Genetic Analysis and QTL Detection on Fiber Traits Using Two Recombinant Inbred Lines and Their Backcross Populations in Upland Cotton. G3 6: 2717–2724.

Shcherban, T. Y., J. Shi, D. M. Durachko, M. J. Guiltinan, S. J. McQueen-Mason et al., 1995 Molecular cloning and sequence analysis of expansins--a highly conserved, multigene family of proteins that mediate cell wall extension in plants. Proc. Natl. Acad. Sci. U. S. A. 92: 9245–9249.

Shen, X., T. Zhang, W. Guo, X. Zhu, and X. Zhang, 2006 Mapping Fiber and Yield QTLs with Main, Epistatic, and QTL × Environment Interaction Effects in Recombinant Inbred Lines of Upland Cotton. Crop Sci. 46: 61–66.

Shikata, M., T. Koyama, N. Mitsuda, and M. Ohme-Takagi, 2009 Arabidopsis SBP-box genes SPL10, SPL11 and SPL2 control morphological change in association with shoot maturation in the reproductive phase. Plant Cell Physiol. 50: 2133–2145.

Shi, Y.-H., S.-W. Zhu, X.-Z. Mao, J.-X. Feng, Y.-M. Qin et al., 2006 Transcriptome profiling, molecular biological, and physiological studies reveal a major role for ethylene in cotton fiber cell elongation. Plant Cell 18: 651–664.

Signor, S. A., and S. V. Nuzhdin, 2018 The Evolution of Gene Expression in cis and trans. Trends Genet. 34: 532–544.

Song, C., W. Li, X. Pei, Y. Liu, Z. Ren et al., 2019 Dissection of the genetic variation and candidate genes of lint percentage by a genome-wide association study in upland cotton. Theor. Appl. Genet. 132: 1991–2002.

Spokevicius, A. V., S. G. Southerton, C. P. MacMillan, D. Qiu, S. Gan et al., 2007 β-tubulin affects cellulose microfibril orientation in plant secondary fibre cell walls: β-tubulin affects MFA. Plant J. 51: 717–726.

Stam, P., 1993 Construction of integrated genetic linkage maps by means of a new computer package: Join Map. Plant J. 3: 739–744.

Steinborn, K., C. Maulbetsch, B. Priester, S. Trautmann, T. Pacher et al., 2002 The Arabidopsis PILZ group genes encode tubulin-folding cofactor orthologs required for cell division but not cell growth. Genes Dev. 16: 959–971.

Taliercio, E. W., and D. Boykin, 2007 Analysis of gene expression in cotton fiber initials. BMC Plant Biol. 7: 22.

Tan, Z., X. Fang, S. Tang, J. Zhang, D. Liu et al., 2015 Genetic map and QTL controlling fiber quality traits in upland cotton (Gossypium hirsutum L.). Euphytica 203: 615–628.

Tang, W., Q. Ji, Y. Huang, Z. Jiang, M. Bao et al., 2013 FAR-RED ELONGATED HYPOCOTYL3 and FAR-RED IMPAIRED RESPONSE1 transcription factors integrate light and abscisic acid signaling in Arabidopsis. Plant Physiol. 163: 857–866.

Tang, S., Z. Teng, T. Zhai, X. Fang, F. Liu et al., 2015 Construction of genetic map and QTL analysis of fiber quality traits for Upland cotton (Gossypium hirsutum L.). Euphytica 201: 195–213.

Tan, Z., Z. Zhang, X. Sun, Q. Li, Y. Sun et al., 2018 Genetic Map Construction and Fiber Quality QTL Mapping Using the CottonSNP80K Array in Upland Cotton. Front. Plant Sci. 9: 225.

Tuttle, J. R., G. Nah, M. V. Duke, D. C. Alexander, X. Guan et al., 2015 Metabolomic and transcriptomic insights into how cotton fiber transitions to secondary wall synthesis, represses lignification, and prolongs elongation. BMC Genomics 16: 477.

Tyagi, P., M. A. Gore, D. T. Bowman, B. T. Campbell, J. A. Udall et al., 2014 Genetic diversity and population structure in the US Upland cotton (Gossypium hirsutum L.). Theor. Appl. Genet. 127: 283–295.

Ulloa, M., S. Saha, J. N. Jenkins, W. R. Meredith Jr, J. C. McCarty Jr et al., 2005 Chromosomal assignment of RFLP linkage groups harboring important QTLs on an intraspecific cotton (Gossypium hirsutum L.) Joinmap. J. Hered. 96: 132–144.

Ungerer, M. C., S. S. Halldorsdottir, J. L. Modliszewski, T. F. C. Mackay, and M. D. Purugganan, 2002 Quantitative trait loci for inflorescence development in Arabidopsis thaliana. Genetics 160: 1133– 1151.

Van der Auwera, G. A., M. O. Carneiro, C. Hartl, R. Poplin, G. del Angel et al., 2013 From FastQ Data to High-Confidence Variant Calls: The Genome Analysis Toolkit Best Practices Pipeline. Current Protocols in Bioinformatics 43: 11.10.1–11.10.33.

VAN Ooijen, J. W., 2011 Multipoint maximum likelihood mapping in a full-sib family of an outbreeding species. Genet. Res. 93: 343–349.

Voorrips, R. E., 2002 MapChart: Software for the Graphical Presentation of Linkage Maps and QTLs. Journal of Heredity 93: 77–78.

Wang, S., C. J. Basten, and Z.-B. Zeng, 2012 Windows QTL Cartographer 2.5. Windows QTL Cartographer.

Wang, B., X. Draye, Z. Zhang, Z. Zhuang, O. L. May et al., 2016a Advanced Backcross Quantitative Trait Locus Analysis of Fiber Elongation in a Cross between Gossypium hirsutum and G. mustelinum. Crop Sci. 56: 1760–1768.

Wang, B., X. Draye, Z. Zhuang, Z. Zhang, M. Liu et al., 2017a QTL analysis of cotton fiber length in advanced backcross populations derived from a cross between Gossypium hirsutum and G. mustelinum. Theor. Appl. Genet. 130: 1297–1308.

Wang, H., C. Huang, H. Guo, X. Li, W. Zhao et al., 2015 QTL Mapping for Fiber and Yield Traits in Upland Cotton under Multiple Environments. PLoS One 10: e0130742.

Wang, F., W. Kong, G. Wong, L. Fu, R. Peng et al., 2016b AtMYB12 regulates flavonoids accumulation and abiotic stress tolerance in transgenic Arabidopsis thaliana. Mol. Genet. Genomics 291: 1545– 1559.

Wang, Y., and J. Li, 2006 Genes controlling plant architecture. Curr. Opin. Biotechnol. 17: 123–129.

Wang, Q. Q., F. Liu, X. S. Chen, X. J. Ma, H. Q. Zeng et al., 2010 Transcriptome profiling of early developing cotton fiber by deep-sequencing reveals significantly differential expression of genes in a fuzzless/lintless mutant. Genomics 96: 369–376.

Wang, M., L. Tu, M. Lin, Z. Lin, P. Wang et al., 2017b Asymmetric subgenome selection and cis-regulatory divergence during cotton domestication. Nat. Genet. 49: 579–587.

Wang, B., Z. Zhuang, Z. Zhang, X. Draye, L.-S. Shuang et al., 2017c Advanced Backcross QTL Analysis of Fiber Strength and Fineness in a Cross between Gossypium hirsutum and G. mustelinum. Front. Plant Sci. 8: 1848.

Wang, F., D. Zhu, X. Huang, S. Li, Y. Gong et al., 2009 Biochemical insights on degradation of Arabidopsis DELLA proteins gained from a cell-free assay system. Plant Cell 21: 2378–2390.

Waters, A. J., I. Makarevitch, J. Noshay, L. T. Burghardt, C. N. Hirsch et al., 2017 Natural variation for gene expression responses to abiotic stress in maize. Plant J. 89: 706–717.

Wendel, J. F., and V. A. Albert, 1992 Phylogenetics of the Cotton Genus (Gossypium): Character-State Weighted Parsimony Analysis of Chloroplast-DNA Restriction Site Data and Its Systematic and Biogeographic Implications. Syst. Bot. 17: 115–143.

Wendel, J. F., and C. E. Grover, 2015 Taxonomy and Evolution of the Cotton Genus, Gossypium, pp. 25– 44 in Cotton, Agronomy Monograph, American Society of Agronomy, Inc., Crop Science Society of America, Inc., and Soil Science Society of America, Inc., Madison, WI.

Wen, J., K. A. Lease, and J. C. Walker, 2004 DVL, a novel class of small polypeptides: overexpression alters Arabidopsis development. Plant J. 37: 668–677.

Wittkopp, P. J., B. K. Haerum, and A. G. Clark, 2004 Evolutionary changes in cis and trans gene regulation. Nature 430: 85–88.

Wu, T. D., and S. Nacu, 2010 Fast and SNP-tolerant detection of complex variants and splicing in short reads. Bioinformatics 26: 873–881.

Wu, T. D., and C. K. Watanabe, 2005 GMAP: a genomic mapping and alignment program for mRNA and EST sequences. Bioinformatics 21: 1859–1875.

Xiao, Y.-H., Q. Yan, H. Ding, M. Luo, L. Hou et al., 2014 Transcriptome and biochemical analyses revealed a detailed proanthocyanidin biosynthesis pathway in brown cotton fiber. PLoS One 9: e86344.

Xiao, Y.-H., Z.-S. Zhang, M.-H. Yin, M. Luo, X.-B. Li et al., 2007 Cotton flavonoid structural genes related to the pigmentation in brown fibers. Biochem. Biophys. Res. Commun. 358: 73–78.

Xiao, G., P. Zhao, and Y. Zhang, 2019 A Pivotal Role of Hormones in Regulating Cotton Fiber Development. Front. Plant Sci. 10: 87.

Xu, X. M., A. Rose, S. Muthuswamy, S. Y. Jeong, S. Venkatakrishnan et al., 2007 NUCLEAR PORE ANCHOR, the Arabidopsis homolog of Tpr/Mlp1/Mlp2/megator, is involved in mRNA export and SUMO homeostasis and affects diverse aspects of plant development. Plant Cell 19: 1537–1548.

Yadav, V., I. Molina, K. Ranathunge, I. Q. Castillo, S. J. Rothstein et al., 2014 ABCG transporters are required for suberin and pollen wall extracellular barriers in Arabidopsis. Plant Cell 26: 3569–3588.

Yam, K. L., and S. E. Papadakis, 2004 A simple digital imaging method for measuring and analyzing color of food surfaces. J. Food Eng. 61: 137–142.

Yoo, M.-J., and J. F. Wendel, 2014 Comparative evolutionary and developmental dynamics of the cotton (Gossypium hirsutum) fiber transcriptome. PLoS Genet. 10: e1004073.

Yu, J., S. Jung, C.-H. Cheng, S. P. Ficklin, T. Lee et al., 2014 CottonGen: a genomics, genetics and breeding database for cotton research. Nucleic Acids Res. 42: D1229–36.

Zeng, Z. B., 1994 Precision mapping of quantitative trait loci. Genetics 136: 1457–1468.

Zeng, Z. B., 1993 Theoretical basis for separation of multiple linked gene effects in mapping quantitative trait loci. Proc. Natl. Acad. Sci. U. S. A. 90: 10972–10976.

Zhang, M., L.-B. Han, W.-Y. Wang, S.-J. Wu, G.-L. Jiao et al., 2017 Overexpression of GhFIM2 propels cotton fiber development by enhancing actin bundle formation. J. Integr. Plant Biol. 59: 531–534.

Zhang, T., Y. Hu, W. Jiang, L. Fang, X. Guan et al., 2015 Sequencing of allotetraploid cotton (Gossypium hirsutum L. acc. TM-1) provides a resource for fiber improvement. Nat. Biotechnol. 33: 531–537.

Zhang, J., R. G. Percy, and J. C. McCarty, 2014 Introgression genetics and breeding between Upland and Pima cotton: a review. Euphytica 198: 1–12.

Zhang, Y., X. F. Wang, Z. K. Li, G. Y. Zhang, and Z. Y. Ma, 2011 Assessing genetic diversity of cotton cultivars using genomic and newly developed expressed sequence tag-derived microsatellite markers. Genet. Mol. Res. 10: 1462–1470.

Zhang, Z.-S., Y.-H. Xiao, M. Luo, X.-B. Li, X.-Y. Luo et al., 2005 Construction of a genetic linkage map and QTL analysis of fiber-related traits in upland cotton (Gossypium hirsutum L.). Euphytica 144: 91–99.

Zhao, Y., H. Wang, W. Chen, Y. Li, H. Gong et al., 2015 Genetic diversity and population structure of elite cotton (Gossypium hirsutum L.) germplasm revealed by SSR markers. Plant Syst. Evol. 301: 327–336.

